# Stress Resistance Screen in a Human Primary Cell Line Identifies Small Molecules that Affect Aging Pathways and Extend *C. elegans’* Lifespan

**DOI:** 10.1101/735415

**Authors:** Peichuan Zhang, Yuying Zhai, James Cregg, Kenny Kean-Hooi Ang, Michelle Arkin, Cynthia Kenyon

## Abstract

Increased resistance to environmental stress at the cellular level is correlated with the longevity of long-lived mutants and wild-animal species. Moreover, in experimental organisms, screens for increased stress resistance have yielded mutants that are long-lived. To find entry points for small molecules that might extend healthy longevity in humans, we screened ∼100,000 small molecules in a human primary-fibroblast cell line and identified a set that increased oxidative-stress resistance. Some of the hits fell into structurally-related chemical groups, suggesting that they may act on common targets. Two small molecules increased *C. elegans’* stress resistance, and at least 9 extended their lifespan by ∼10-50%. Thus, screening for increased stress resistance in human cells can enrich for compounds with promising pro-longevity effects. Further characterization of these compounds, including a chalcone that promoted stress resistance independently of *NRF2*, may elucidate new ways to extend healthy human lifespan.

## Introduction

In animals, mutations in nutrient, energy and stress-sensing genes, such as IGF-1-axis and mTOR-signaling genes, extend youthfulness and lifespan and counter age-related disease (Bartke, 2011; Fontana et al., 2010; Kenyon, 2010a). These genes are members of large networks with multiple components, some of which could potentially serve as targets for pharmacological interventions to increase healthy lifespan. In fact, consistent with genetic perturbations, small molecules that inhibit mTOR (rapamycin) or activate AMP kinase (metformin) have been reported to prolong lifespan in several different species (Ding et al., 2017), albeit modestly in the case of metformin in mice (Martin-Montalvo et al., 2013). Some of these small molecules have been used in humans to treat diabetes and cancer, two diseases that afflict the aging population. Together these findings suggest that interventions targeting components of this network could extend the healthy life of humans.

In addition to targeting human orthologs of proteins known to influence longevity in animals, an alternative approach might be to identify promising molecules in phenotypic screens for cellular correlates of animal longevity. One such correlate is resistance to environmental stress. Many long-lived mutants and their cells are resistant to multiple types of stressors (Miller, 2009). In addition, cells from long-lived wild animals, such as brown bats and naked mole rats, are resistant to oxidizing radicals, heavy metals and DNA-damaging agents (Harper et al., 2007; Lewis et al., 2012; Salmon et al., 2008). Third, forkhead box O (FOXO) proteins (such as FOXO3) and nuclear factor, erythroid 2 like 2 (NFE2L2, aka NRF2), key regulators of stress responses, can promote longevity in several species, including worms (Bishop and Guarente, 2007; Kenyon et al., 1993; Libina et al., 2003; Ogg et al., 1997; Tullet et al., 2008), flies (Giannakou et al., 2004; Hwangbo et al., 2004; Sykiotis and Bohmann, 2008), and likely, mice (Leiser and Miller, 2010; Shimokawa et al., 2015; Steinbaugh et al., 2012). In humans, exceptional longevity-associated *FOXO3A* polymorphisms have been identified in multiple cohorts (Joshi et al., 2017; Kenyon, 2010a; Kolovou et al., 2017), and NRF2 activation and mTOR inhibition have been shown to delay senescence in human fibroblasts (Kapeta et al., 2010; Walters et al., 2016). As proof of concept, genetic screens for organismal stress resistance in yeast, worms and plants have enriched for mutants that live long (Cao et al., 2003; de Castro et al., 2004; Kennedy et al., 1995; Kim and Sun, 2007). Conversely, by screening a library of compounds with known mammalian pharmacological targets, Ye and coworkers identified 60 that promoted longevity of *C. elegans*, and of these, 33 increased worms’ resistance to oxidative stress (Ye et al., 2014). Several lines of evidence indicate that increased oxidative-stress resistance may not be the cause of extended longevity (Perez et al., 2009; Slack et al., 2011). However, these findings show that selection strategies for stress resistance can be used to enrich for longevity regulators.

To this end, we carried out a high-throughput screen for small molecules that enhance resistance to hydrogen peroxide-induced oxidative stress in a human primary cell line. We further characterized our hit molecules in assays related to known longevity pathways and found certain molecules activated targets of NRF2/SKN-1. We also found that some molecules could extend the lifespan of *C. elegans*.

## Results

### Small-Molecule Screen for Oxidative-Stress Resistance

Primary skin fibroblasts from long-lived mouse mutants and long-lived animal species exhibit cellular resistance to the stressors hydrogen peroxide, cadmium and methyl methanesulfonate (MMS) (Harper et al., 2007; Harper et al., 2011; Salmon et al., 2005). For our screen, we used the WI-38 human primary-fibroblast cell line. Tumor cells were avoided because conditions that extend animal lifespan generally inhibit tumor cell growth (Cheng et al., 2014; Kalaany and Sabatini, 2009; Pinkston et al., 2006). We note that not all types of cells derived from long-lived and stress-resistant animals are stress-resistant in culture. Some, such as hepatocytes, are more likely to undergo apoptosis (Kennedy et al., 2003). Likewise, human mammary epithelial cells incubated in IGF-1-deficient serum from humans defective in growth-hormone receptor are sensitized to die when exposed to hydrogen peroxide (Guevara-Aguirre et al., 2011).

In the screen, we looked for small molecules that increased cellular resistance to hydrogen peroxide (Figure 1). The screening flow is summarized in Figure 1A and detailed in Materials & Methods. Briefly, we screened a compound collection that contains 104,121 structurally diverse small molecules selected to maximize the coverage of chemical space. In our primary screen, we assessed cell viability by measuring ATP content. We selected a 3-hour treatment with 700 μM H_2_O_2_, as, importantly, small interfering RNAs (siRNAs) against the insulin/IGF-1 signal-transducing gene *AKT1* and the NRF2 inhibitor gene *KEAP1* both increased stress resistance under these conditions (Supplemental Figure 1), suggesting that we could recover the types of perturbations that are known to increase lifespan in animals. Altogether, we identified 209 small molecules (0.2% of molecules tested, at 10 μM) that produced signals at least 2.5-fold greater than the DMSO-treated negative control (Supplemental Table 1). We retested these candidates at six different concentrations (0.6 μM to 20 μM) and confirmed 127 that consistently produced protective effects against H_2_O_2_ at one or more of these doses (data not shown).

**Figure 1.**
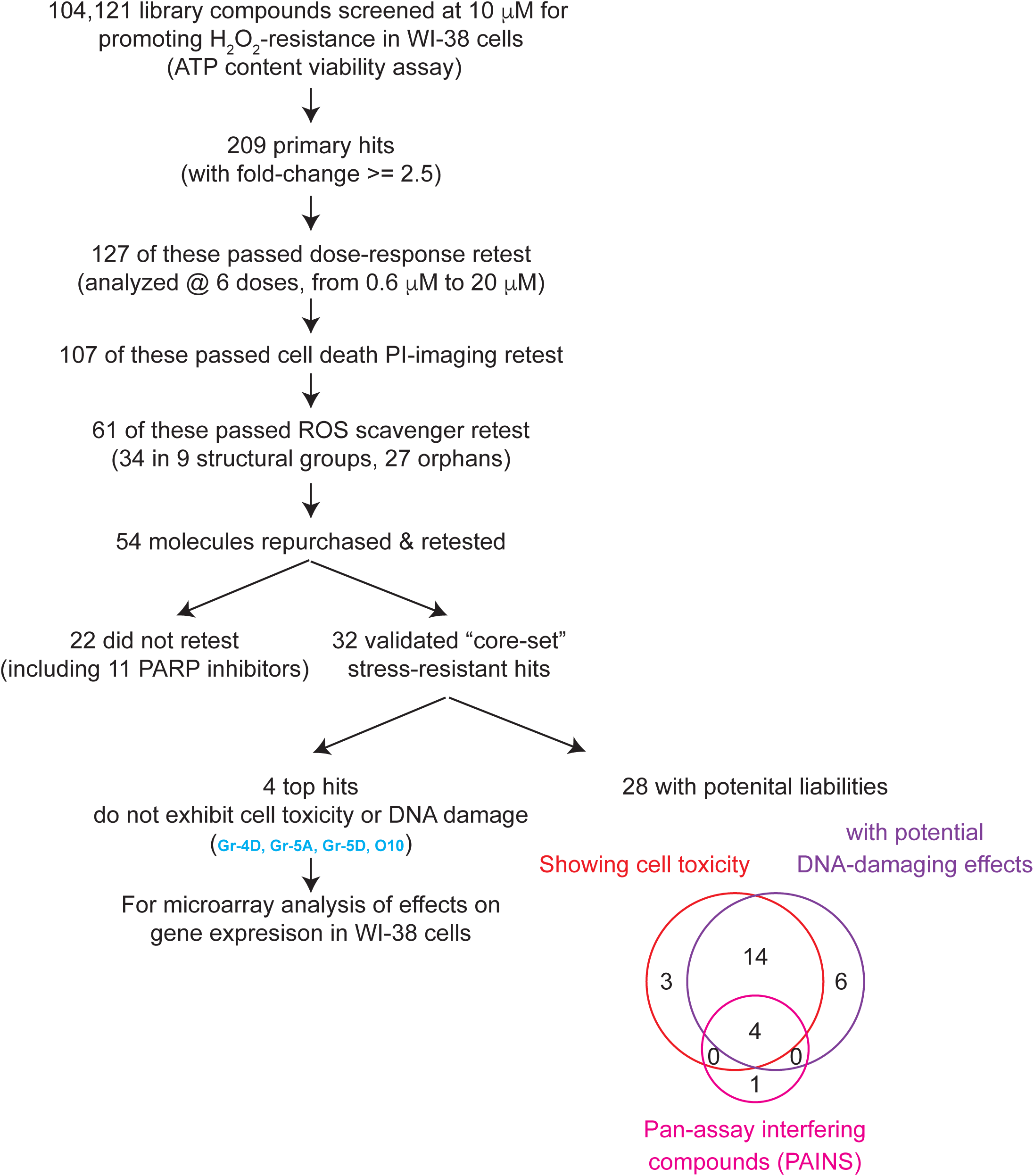
Summary of the small-molecule screen hits. A cell viability assay was performed for each of 104,121 compounds analyzed, by measuring ATP content, for WI-38 cells that were pre-incubated with 10 μM small molecules overnight and then treated with H_2_O_2_ for 3 hours. 209 primary hits were identified and subjected to a series of retests to eliminate false positives and ROS scavengers to yield the top 61 selected hits. Of these, a core set of 32 small molecules were validated and further analyzed in cells and in worms. Many molecules showed cell toxicity and potential DNA-damaging effects, and 4 top non-toxic molecules showing no liabilities were analyzed for their effects on gene expression in WI-38 cells.

To exclude molecules that did not increase stress resistance but instead somehow elevated cellular ATP levels (Thorne et al., 2010), we also carried out a secondary imaging assay for cell viability. Using propidium iodide (PI), a cell non-permeable dye that stains DNA in late-apoptotic/necrotic cells when membrane integrity is lost, we identified 107 compounds that reduced the percentage of dead cells following H_2_O_2_ stress. Predictably, among the molecules that increased ATP levels but produced little or no protective effect in PI-imaging assays were several inhibitors of poly ADP-ribose polymerase (PARP) (see Supplemental Text). PARP consumes ATP to repair DNA damage; thus, its inhibition increases ATP levels without protecting cells from oxidative stress.

Among these 107 hits were 32 potential derivatives of 8-hydroxyquinoline (8-HQ), a well-known reactive oxygen species (ROS) scavenger that can protect cells from H_2_O_2_ stress (Wang et al., 2010). We excluded additional such compounds using an *in vitro* ROS scavenger assay (Supplemental Figure 3), leaving 61 primary hits (Figure 2; Supplemental Table 1). Of these, 27 had unique chemical structures (orphans) whereas 34 fell into one of nine structural classes containing either two or more members (Gr1 to Gr9), suggesting that they may act on common targets to protect cells from H_2_O_2_. These structures are shown in Supplemental Table 1.

**Figure 2.**
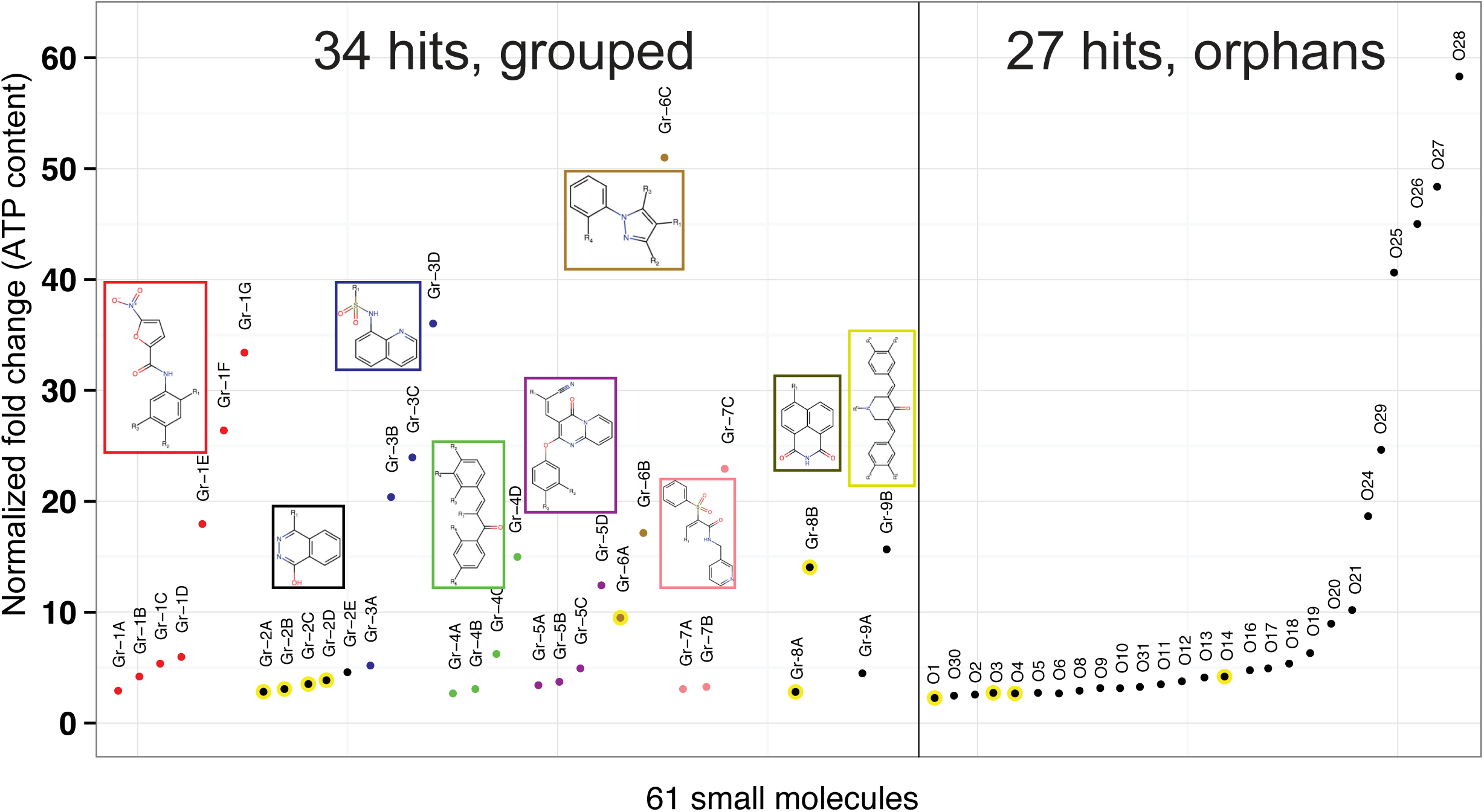
Normalized fold changes (ATP content) of H_2_O_2_-treated WI-38 cells, which were pre-incubated with each of the top 61 hit compounds that passed the initial selection criteria (see Figure 1). 34 molecules fell into one of nine groups (Gr-1 to 9, shown in different colors – core structures are shown) (see Supplemental Table 1 for structures). Each group comprised two or more members that share a similar core structure. In addition, we identified 27 orphan compounds. Yellow circles: PARP inhibitors (see Supplemental Figure 8).

We were able to obtain a fresh batch of 54 of the 61 compounds and verified their molecular masses, with 3 exceptions. However, we noted that, among these 51, several batches of compounds that we continued to analyze (Gr-5A, Gr-5B, Gr-6B, Gr-6C and O27) contained a second LC-MS peak, likely a modified species. We first verified, by performing an *in vitro* Amplex Red assay, that these molecules did not quench H2O2 (Supplemental Figure 4), excluding this as a possible explanation for their protective effects. We then confirmed in multiple independent experiments, by measuring ATP content and performing additional PI-imaging, that 32 molecules (excluding all the PARP inhibitors, some of which scored borderline in the PI-imaging assays) reproducibly protected WI-38 cells during H_2_O_2_ treatment (Supplemental Table 1 & 2). Hence these 32 small molecules became our “core set” of hits for further characterization (Table 1).

**Table 1.** Summary of core-set small molecule hits by their phenotypic assay scores. 32 core-set hits, including 22 that belong to 6 structural groups and 10 orphans, were analyzed in multiple longevity-correlated assays and their behaviors are shown in this table. Their relative effects in each assay are indicated (also see Supplemental Table 1). Asterisk (*): molecules with signature structures of pan-assay interference (PAIN) compounds. Four top hit compounds showing no obvious cell toxicity are highlighted in blue. N.D. not determined.

| ID | Cell-based assays |  |  |  |  |  | Worm-based assays |  | Liabilities |  |
| --- | --- | --- | --- | --- | --- | --- | --- | --- | --- | --- |
|  | H<sub>2</sub>O<sub>2</sub>-resistant | Cd-resistant | Potential FOXO3 Activation (by qPCR assay) | Potential NRF2 Activation (by qPCR assay) | mTOR down-regulation | Protection from Huntington's poly-Q toxicity | H<sub>2</sub>O<sub>2</sub>-resistance in worms | Lifespan-extending scores in worms | Potential DNA-damaging | Potential toxicity based on ATP and confluency assays |
| **Top, non-toxic molecules (4)** |  |  |  |  |  |  |  |  |  |  |
| Gr-4D | yes | ++ | no | ++ | no | no | + | +++ | no | no |
| Gr-5A | yes | ++ | no | + | no | no | + | no | no | no |
| Gr-5D | yes | no | no | + | no | no | no | no | no | no |
| O10 | yes | ++ | no | no | no | no | no | no | no | no |
| **Others with potential liability (PAINS, indicated) (28)** |  |  |  |  |  |  |  |  |  |  |
| Gr-1A | yes | + | no | no | no | + | + | no | + | no |
| Gr-1B | yes | ++ | no | no | no | no | + | + | + or no | no |
| Gr-1C | yes | ++ | no | no | no | + | no | + | + | no |
| Gr-1D | yes | ++ | no | no | no | no | no | + | + | no |
| Gr-1E | yes | ++ | no | no | no | no | no | ++ | + | + |
| Gr-1F | yes | ++ | no | no | no | no | no | + | + | + |
| Gr-1G | yes | ++ | no | no | no | no | no | + | + | + |
| Gr-3A | yes | ++ | no | no | no | no | no | ++ | + | no |
| Gr-3B | yes | no | no | no | no | no | + | ++ | + | + |
| Gr-3C | yes | no | no | no | no | no | no | ++ | + | + |
| Gr-4A | yes | ++ | no | ++ | no | no | no | no | + or no | + |
| Gr-4B | yes | ++ | no | ++ | no | no | no | + | no | + |
| Gr-4C | yes | no | no | ++ | no | no | no | no | no | + |
| Gr-5B | yes | no | no | + | no | no | no | no | + or no | no |
| Gr-6B | yes | ++ | no | + | no | + | no | no | + or no | + |
| Gr-7A | yes | N.D. | + | no | + | no | ++ | ++ | + | + |
| Gr-7B | yes | N.D. | no | no | + | no | + | no | + | no |
| Gr-7C | yes | no | + | + | + | + | no | no | + or no | + |
| O11 | yes | ++ | no | no | no | no | + | + | + or no | + |
| O13 | yes | + | no | + | no | no | + | +++ | + or no | + |
| O18 | yes | no | no | + | no | no | no | + | + or no | + |
| O20 | yes | ++ | no | + | no | no | no | + | + or no | no |
| O27 | yes | ++ | no | no | + | no | no | — | + | + |
| O6 (* non-toxic) | yes | ++ | no | + | no | no | no | + | no | N.D. |
| O21 (*) | yes | no | no | no | no | no | no | no | + | + |
| Gr-6C (*) | yes | no | no | no | no | no | no | ++ | + | + |
| Gr-9A (*) | yes | N.D. | no | + | no | no | ++ | + | + or no | + |
| Gr-9B (*) | yes | N.D. | + | no | no | no | + | ++ | + or no | + |

A second cellular stress that correlates well with organismal longevity is resistance to cadmium, which also produces ROS (Harper et al., 2007). Therefore, we analyzed the ability of our core-set small molecules to protect WI-38 cells from cadmium, using the same ATP assay. At least 17 molecules increased resistance to cadmium as well as H_2_O_2_ (Supplemental Table 3). Furthermore, 22 validated hits also protected human primary dermal fibroblasts (isolated from the skin of multiple donors) from H_2_O_2_ (2 trials, Supplemental Table 2), demonstrating that their protective capacity was not limited to the WI-38 cell line. We also asked whether any molecules could increase resistance of cells to the DNA-damaging agent MMS, but none of our 32 core-set hits scored positively in this assay.

Finally, we sought to eliminate small molecules that had obvious toxicities at the lowest efficacious dose. To this end, we analyzed long-term effects of the core-set molecules on ATP levels and cell confluency in culture (see Supplemental Text; Supplemental Figure 5 & Supplemental Table 4), and we also tested their ability to induce proliferation arrest or DNA damage-associated markers in the absence of H_2_O_2_ (Supplemental Table 4). We found that 24 of our 32 core-set small molecules elicited signs of DNA damage; 21 of these also produced cell toxicity (assayed either by effects on ATP content or cell confluency) (Table 1). However, five molecules exhibited none of these potential liabilities: Gr-4D, Gr-5A, Gr-5D, O6 and O10 (see Supplemental Figure 5).

Electrophilic “pan assay interference compounds” (PAINS), due to their reactive nature, can produce non-specific effects in a chemical screen (Baell and Walters, 2014; Baell and Holloway, 2010) (See Supplemental Text for more discussions). Of our 32 core hits, one non-toxic compound, O6, fit the criteria of a PAIN, as did 4 of the toxic compounds. Because of their relatively attractive features for drug development, the 4 non-toxic, non-PAIN molecules (Gr-4D, Gr-5A, Gr-5D and O10) were investigated most extensively in this study. However, because the toxicity we observed with the other compounds could potentially be due to off-target effects, and, interestingly, because low levels of some toxic agents (such as paraquat) have been shown to induce protective responses that extend life, we ran several additional tests on all 32 compounds.

### Effects of Hit Compounds on Gene Expression

To gain insight into potential mechanisms by which the small molecules promoted oxidative stress-resistance, we performed RNA-seq analysis on our most promising hit, the chalcone Gr-D4, which extended the lifespan of *C. elegans* up to 50% (described below; see Figure 4 & Supplemental Table 8) and carried out microarray analysis of additional hits.

**Figure 3.**
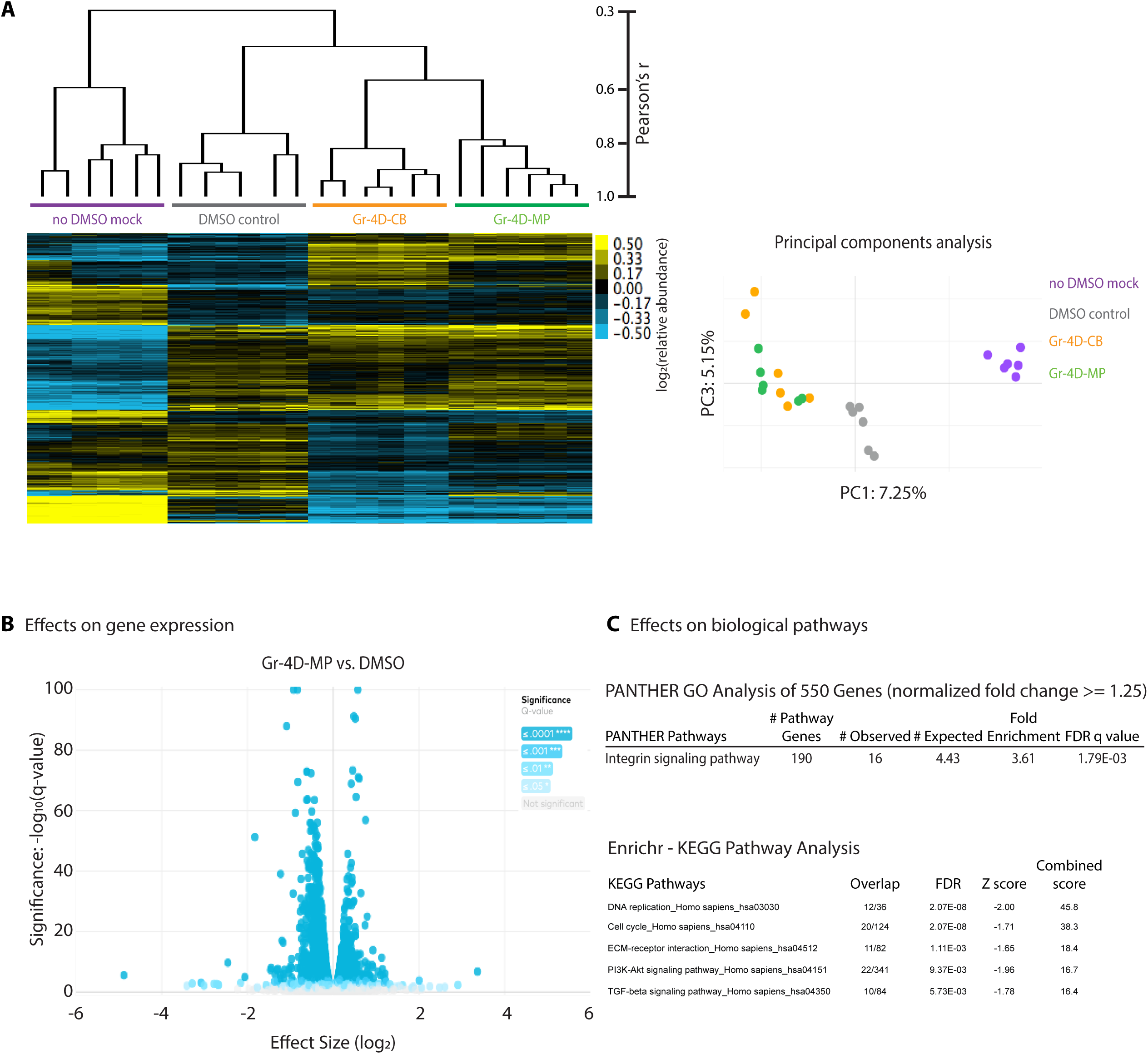
Effects of Gr-4D on gene expression of WI-38 cells. A) Pearson’s correlation between global transcriptional profiles for cells treated with Gr-4D. Shown is the normalized gene expression matrices (transcript per million, TPM values) of the 8,000 genes detected in every RNA-seq sample, grouped by unsupervised clustering using Pearson correlation coefficient as a distance metric. Note that most DMSO (0.1%) controls were clustered together on the tree, as were the no-DMSO mock controls or samples treated with Gr-4D (2.5 μM) from two different vendors (CB, ChemBridge; MP, MolPort). This correlation was observed also in the principle components analysis (PCA), by projecting two major components (PC1 and PC3), which explained 7.25% and 5.15% variation of global gene expression, respectively B) Volcano plot showing effects of Gr-4D on gene expression. Shown are the normalized expression level (X axis) and FDR-adjusted significance (Y axis) for one Gr-4D (from MolPort). Each dot represents a gene that was differentially expressed in WI-38 cells upon treatment with Gr-4D (see Supplemental Table 6A). C) Pathway analysis indicating effects of Gr-4D on certain pathways. Shown are the top pathways affected (by PANTHER analysis of 550 significant genes, with FDR-adjust q value of overlap with the known pathway genes < 0.05; or by Enrichr-KEGG pathway analysis, with a ranking score combining both FDR and z score > 10) (see Supplemental Table 6A).

**Figure 4.**
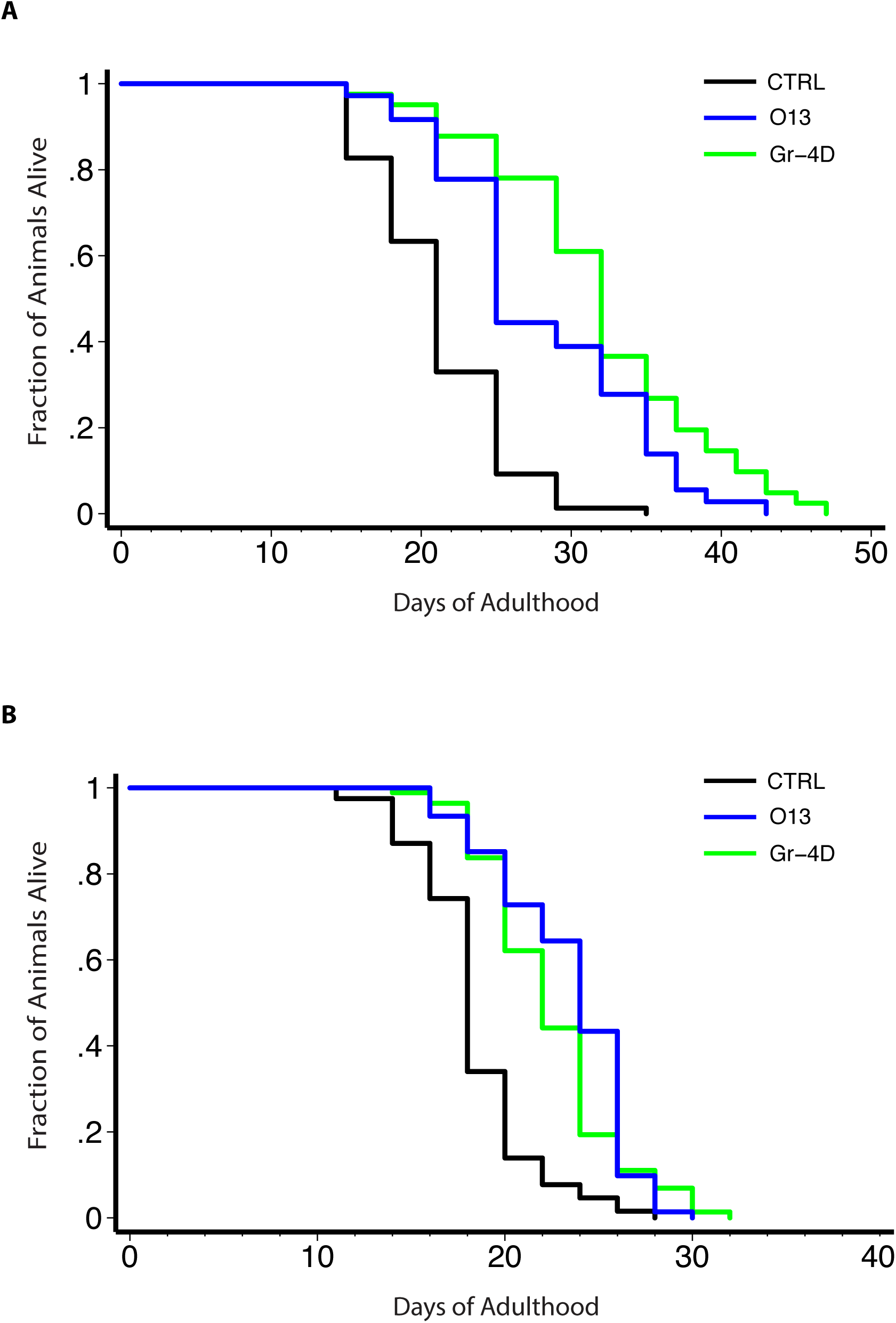
Extension of *C. elegans’* lifespan by small molecules. Small molecules were analyzed for their ability to extend the lifespan of *C. elegans*. Two small molecules (Gr-4D, without cell toxicity; O13, with cell toxicity) that consistently extended lifespan in multiple independent assays are shown. A) Wild-type animals, grown in liquid, with FuDR to block progeny production, 20°C. Control, 21.2±0.5 (mean ± SEM in days), n = 77/81 (observed/total); Gr-4D-treated, 31.9±1.1 (50.5% increase), n = 41/41, *P* < 0.0001 (log-rank test); O13-treated, 28.1±1.1 (32.5% increase), n = 36/36, *P* < 0.0001. B) Temperature-sensitive sterile mutant animals, grown on plate, without FuDR, 25°C then shifted to room temperature (∼22°C) as adults. Control, 18.4±0.4, n = 67/80; Gr-4D-treated, 22.5±0.4 (22.2% increase), n = 74/95, *P* < 0.0001 (log-rank test); O13-treated, 23.4±0.4 (27.2% increase), n = 72/76, *P* < 0.0001 (see Supplemental Table 8 for details).

Unsupervised hierarchical clustering of the Gr-4D RNA-seq data (which considers global expression variations) showed that, as expected, biological repeats were closely correlated for each condition (Figure 3A). Similarly, this correlation was observed for controls or Gr-4D-treated samples in principle components analysis (Figure 3A). Gr-4D caused rather small changes to most genes. We identified 115 genes whose expression was altered by at least 1.5-fold (vs. DMSO control, false discovery rate < 0.05) consistently in cells treated with Gr-4D molecules from two different vendors (Figure 3B & Supplemental Table 5A). Analysis of these genes did not suggest an obvious enrichment of genes involved in specific biological pathways. When we expanded our analysis to 550 significant genes whose expression showed a modest change (>= 1.25-fold), we observed an enrichment of genes involved in integrin signaling, including multiple collagen-encoding genes whose expression was down-regulated, suggesting an attenuation of this pathway (Figure 3C & Supplemental Table 6A). Further analysis using the Enrichr suite, which integrates multiple types of pathway analysis tools, suggested a significant enrichment of genes involved in DNA replication, cell cycle regulation, PI3K/AKT and TGF-beta signaling (Figure 3C & Supplemental Table 6).

Among the 550 genes modestly changed upon Gr-4D, we did not observe a strong signature for the longevity-associated gene *FOXO3*, but we did observe a modest induction of a number of NRF2 target genes, including nuclear receptor *NR0B1*, NAD(P)H dehydrogenase *NQO1*, aldo-keto reductases *AKR1B10* and *AKR1C1* (Supplemental Table 5A). This was noteworthy, because NRF2/SKN-1 is a key regulator of oxidative stress and xenobiotic phase 2 detoxification, and it can extend animal lifespan (Sykiotis and Bohmann, 2008; Tullet et al., 2008). We also asked whether Gr-4D’s expression profile resembled any signatures identified under other conditions in the iLINCs database, including gene knockdowns and small-molecule perturbations. Interestingly, *KEAP1* knockdowns, which activate NRF2, scored at the top among the genetic perturbations that produced profiles significantly resembling Gr-4D’s signature (Supplemental Table 6). These results suggested that Gr-4D could elicit a modest activation of NRF2, consistent with its identity as a chalcone (see Properties of Small Molecules), compound known to activate NRF2.

We also noted that inhibitors of heat shock protein 90 (HSP90) and histone deacetylases (HDACs), as well as inhibitors of the proteasome, were among the top chemical perturbations that produced gene expression signatures showing very strong and positive correlations to Gr-4D’s (Supplemental Table 6A). HSP90 is known to form a complex with arylhydrocarbon receptor (AhR), a ligand-activated transcription factor that regulates genes such as xenobiotic metabolizing enzymes (Prodromou, 2016). Dissociation of HSP90 upon xenobiotic stimulation could facilitate activation of AhR and enhance NRF2 activity in response to oxidative and xenobiotic stress (Tan and Wahli, 2014). Interestingly, HDAC inhibitors, which could cause hyperacetylation and inhibition of HSP90 (Kovacs et al., 2005), protected neurons against oxidative stress-induced cell death *in vitro* and *in vivo* (Langley et al., 2005). Finally, the proteasome inhibitor MG-132 induced nuclear translocation of NRF2 in human vascular endothelial cells (Sahni et al., 2008); and Nrf2-dependent induction of proteasome and immunoproteasome promoted adaptation to oxidative stress in mouse embryonic fibroblasts (Pickering et al., 2012). Consistent with these findings, we observed that siRNAi knockdown of several proteasome subunits enhanced resistance of WI-38 cells to H_2_O_2_ (Zhang and Chen et al, unpublished).

We also analyzed gene expression by microarray analysis of WI-38 cells treated with 3 other non-toxic molecules (Gr-5A, Gr-5D and O10). For comparison, we included other members of the structural group 4, which produced long-term toxicity. We also included the other group-5 molecule Gr-5B, which, despite its modest induction of TP53BP1, did not appear to exert obvious cell toxicity.

The toxic molecules produced numerous gene-expression changes, but, like Gr-4D, the non-toxic hit compounds induced relatively few significant gene changes (typically less than 500) in the absence of H_2_O_2_ [using Statistical Analysis of Microarrays (SAM) (Tusher et al., 2001), which focuses on highly affected changes] (Supplemental Table 5B). Unsupervised hierarchical clustering of the global gene expression profile showed that biological repeats were also closely correlated, as were the multiple rapamycin-treatment controls (Supplemental Figure 7). Similarly, profiles for the toxic group-4 members also resembled one another and formed a distinctive cluster. Unlike with SAM, using this method, we found that group-5 molecules (Gr-5B & Gr-5D) that share a core structure were loosely correlated with each other. None of the other compounds clustered together.

We performed SAM to identify genes that showed significant changes for each condition (Supplemental Table 5B), and further analyzed their effects on biological systems using pathway analysis tools. Our pathway analysis indicated that, for example, the non-toxic Gr-5A molecule affected genes involved in prostaglandin biosynthesis (Supplemental Table 6). The other group-5 molecule Gr-5D, despite its structural similarity to Gr-5A, affected genes involved in response to DNA damage and stress, as suggested by the down-regulation of multiple genes encoding histones and the up-regulation of genes encoding heat shock proteins. The orphan molecule O10 appeared to induce the integrated stress response (ISR) mediated by ATF4 and PERK, as suggested by the up-regulation of the ISR regulator *CHOP*/*DDIT3*, asparagine synthetase *ASNS* and insulin like growth factor binding protein 1 *IGFBP1*, which is known to be induced by ATF4 upon ER stress (Marchand et al., 2006). Interestingly, ATF4 is up-regulated in long-lived mutant mice (Li et al., 2014). Finally, gene changes induced by the toxic group-4 molecules were strongly and consistently enriched within the NRF2-mediated oxidative stress-response pathway, as suggested by the induction of multiple anti-oxidant and xenobiotic stress response genes (Supplemental Table 5B & 6B).

### Effects of Hit Compounds on Known Longevity Pathways and Protein Aggregation

In addition to analyzing gene expression in cells treated with our 4 best stress resistance-inducing molecules, we also examined all 32 hit compounds from our screen in cellular assays relevant to the biology of aging. Specifically, we asked whether they could activate proteins known to extend lifespan, inhibit mTOR signaling, and/or attenuate toxic protein aggregation.

#### NRF2

In our microarray experiments, a clear signature of NRF2 activity, detected using pathway analysis, was observed only for toxic compounds. To address this issue more broadly, we treated cells with all 32 core-set small molecules and used RT-qPCR to analyze the expression of several canonical NRF2-regulated genes, including *HMOX1* [heme oxygenase (decycling) 1; an anti-oxidant], *NQO1* [NAD(P)H dehydrogenase, quinone 1; a phase 2 detoxification enzyme], *GCLC* (glutamate– cysteine ligase, catalytic; a glutathione-synthesis enzyme) and *GSTM1* (glutathione S transferase) (Hur and Gray, 2011; Suzuki et al., 2013).

Of the 4 best hits described above (Gr-4D, Gr-5A, Gr-5D and O10), three (Gr-4D, Gr-5A and Gr-5D) induced the expression of two or more NRF2-regulated genes at least modestly (by more than 1.5-fold; Supplemental Table 1), suggesting potential NRF2 activation. In addition to these 3 non-toxic compounds, 11 of the remaining 28 toxic or non-specific compounds also appeared to activate NRF2. Of these, Gr-9A induced *HMOX1* by almost 50 fold in WI-38 cells, and this compound also protected *C. elegans* from hydrogen peroxide (in 2 of 2 trials) (Supplemental Table 8). Consistent with the microarray analysis (Supplemental Table 5B), we also noted that all the toxic group-4 molecules resulted in significant induction of NRF2 target genes. These data suggested enrichment for NRF2-activating small molecules in our screen, which is expected, as we selected for agents that could promote oxidative stress-resistance.

NRF2 activation could be one possible mechanism by which our small molecules promote oxidative-stress resistance. In this case, we would expect that *NRF2* to be required for the increased resistance. To address this question, we knocked down *NRF2* by more than 95% in WI-38 cells via siRNA transfection (Supplemental Figure 6A). Compared with *NRF2(+)* cells, *NRF2*-deficent cells showed a high level of propidium iodide staining, and the fraction of PI-positive cells increased further upon treatment with Gr-4D in a dose-dependent manner even in the absence of H_2_O_2_, likely reflecting increased xenobiotic sensitivity upon the loss of *NRF2* (Supplemental Figure 6B). To compensate for this sensitivity to oxidative stress, we treated the *NRF2-* deficient cells with a lower dose of H_2_O_2_ (500uM instead of 700uM), finding that Gr-4D could protect *NRF2(+)* cells under this condition. As expected, compared with *NRF2(+)* cells, *NRF2*-deficient cells were more prone to die upon treatment with H_2_O_2_. To our surprise, Gr-4D (at 3 different doses) was still quite protective, significantly reducing the fraction of PI-positive cells (Supplemental Figure 6B). Thus, *NRF2* was not required for this chalcone to promote oxidative-stress resistance. Gr-4D must act, at least in part, through a different, *NRF2*-independent mechanism.

#### FOXO3

Many perturbations that increase lifespan and stress resistance in animals do so in a *FOXO*-dependent fashion (Kenyon, 2010b). None of our “four-best” compounds scored positively for potential FOXO3A activation. However, 3 toxic compounds, including 2 group-7 molecules, could up-regulate the expression (by more than 1.5-fold) of at least 2 of the 5 FOXO3A-regulated genes analyzed: *SOD2* (superoxide dismutase), *GADD45A* (cell cycle regulator), *CAT* (catalase), *DDB1* (damage-specific DNA binding protein) and *TXNIP* (thioredoxin interacting protein) (Supplemental Table 1).

#### Sestrin

Sestrin genes are regulated by FOXO3 (Nogueira et al., 2008), NRF2 (Shin et al., 2012) and p53 (Budanov et al., 2004). Through direct effects on anti-oxidant peroxiredoxins and through the AMPK and mTOR pathways, sestrins can suppress ROS production and protect cells from oxidative stress, transformation, and genomic instability (Budanov and Karin, 2008). Many studies have indicated that sestrins could be pro-longevity factors. RNAi inhibition of the sestrin gene *sesn-1* has been shown to shorten lifespan, while its overexpression promotes longevity of *C. elegans* (Yang et al., 2013). Loss of *Drosophila dSesn* has been shown to lead to age-associated pathologies, including fat accumulation, mitochondrial dysfunction, muscle degeneration, and cardiac malfunction, which could be blocked by pharmacological activation of AMPK or inhibition of TOR (Lee et al., 2010). Likewise, sestrin deficiencies in mice exacerbated obesity-induced diabetic conditions (Lee et al., 2013; Lee et al., 2012). Furthermore, sestrins have been shown to activate Nrf2 and, by promoting p62-dependent autophagic degradation of Keap1, prevent oxidative damage in the liver of mice (Bae et al., 2013). More recently, the Sabatini group showed that sestrin 2 is a leucine sensor, and that leucine can disrupt sestrin-GATOR2 interaction and result in mTORC1 activation (Wolfson et al., 2016). In our study, of the 32 core-set hits, at least 9 molecules were found to induce the expression (more than 1.5-fold) of *SESN1* (Supplemental Table 1).

Among the four non-toxic compounds, only Gr-4D scored positively for *SESN1* induction. In addition to Gr-4D, 8 of the 28 compounds with liabilities also induced *SESN1*. We noted that these compounds with liabilities, but not Gr-4D, also scored positively for the DNA damage-markers *γ*H2A.X and TP53BP1, and genotoxic stress previously has been shown to up-regulate sestrins through the induction of p53 (Budanov and Karin, 2008). Interestingly, both p53 and Nrf2 are highly expressed in the long-lived, stress-resistant naked mole rat and may contribute to its longevity, and at least in the case of p53, possibly also reduce its risk of cancer (Lewis et al., 2012). In this regard, sestrin-inducing molecules could be useful for promoting healthy aging.

#### mTOR inhibition

Mammalian target of rapamycin (mTOR) is a crucial regulator of cell growth and metabolism and has been implicated in aging and many diseases, including cancer, diabetes and neurological diseases (Dazert and Hall, 2011; Zoncu et al., 2011). This connection has stimulated interest in developing novel mTORC1 inhibitors (Benjamin et al., 2011). In addition to rapamycin and its analogs, the anti-diabetes drug metformin (Kalender et al., 2010), as well as other AMPK activators such as 2-deoxy-D-glucose (Inoki et al., 2003) and AICAR (Shaw et al., 2004), can also inhibit mTORC1.

We examined the effects of our 32 core-set molecules (at three doses, 5 μM, 10 μM and 20 μM) on the phosphorylation status of ribosomal protein S6 (RPS6), a target of mTORC1, and asked whether they could potentially inhibit mTOR. Rapamycin, as a control, reduced the normalized ratio of phosphorylated RPS6 (p-RPS6) by ∼95%.

None of the non-toxic compounds appeared to inhibit mTOR. However, we found that 4 toxic molecules (all 3 members of the 7^th^ structural group and O27) reproducibly reduced the normalized ratio of p-RPS6 (∼30% to 60%, Supplemental Table 7); note that O27 also modestly induced sestrin, a known inhibitor of mTORC1.

#### Poly-Q toxicity attenuation

Protein aggregation is a key feature of many age-related neurodegenerative diseases (Caughey and Lansbury, 2003; Ross and Poirier, 2004). A number of cell lines have been established to study these diseases at the cellular level (Schlachetzki et al., 2013). Among these, PC12 rat pheochromocytoma cells that stably express GFP fused to the poly(Q) tract (exon 1) of mutant human huntingtin gene *HTT* (Q103-Htt-EGFP), have been used to study Huntington’s disease (Aiken et al., 2004). In this cell line, induced expression of poly(Q)103-Htt-EGFP causes the formation of aggregates and exerts substantial cell toxicity. Previously, a medium-throughput small-molecule screen led to the identification of several compounds that could protect cells from toxic Huntington’s aggregates (Aiken et al., 2004), including caspase inhibitors (e.g., Z-VAD-FMK) and anti-inflammatory cannabinoids that have been proposed as therapies in Alzheimer’s and Parkinson’s disease (Aso and Ferrer, 2014; Booz, 2011; Rohn, 2010). We analyzed our repurchased molecules (at 10 μM) in these cells, and then retested several candidates at multiple doses (2.5 μM, 5 μM, 10 μM and 20 μM).

Among the non-toxic compounds, we did not find any that scored positively in this assay. Instead, we found that four toxic molecules (Gr-1A, Gr-1C, Gr-6B and Gr-7C) appeared to increase the ATP content of PC12 cells upon the induction of poly(Q)103-Htt-EGFP aggregates (Supplemental Figure 9). PubChem query indicated that Gr-3B (12 μM) and Gr-4D (5 μM) were identified by another screen and confirmed to be protective against Huntington’s toxicity in the same PC12 cell line (Supplemental Table 9). The basis of this discrepancy is not clear.

### Effects of Hit Compounds on C. elegans

#### Hydrogen peroxide resistance

We analyzed our 32 core hits and found that at least 2 small molecules (Gr-7A & Gr-9A, in 2 trials) could reproducibly increase *C. elegans’* resistance to a lethal dose of hydrogen peroxide (Supplemental Table 1 & 8). Both of these molecules caused toxicity in cultured mammalian cells. We noted that compounds that scored negatively in *C. elegans* are not necessarily uninteresting. As these molecules were identified in a human cell-based screen, their bioavailability or target engagement could be different in worms.

#### Lifespan extension

Because most conditions that extend *C. elegans’* lifespan increase stress resistance, we analyzed how our small molecules affected lifespan. To ensure robustness, we used several different assay conditions (culturing in liquid, on OP50 bacteria-seeded agar plates, on live or UV-irradiated bacteria, and using FuDR-treated or genetically-induced sterile animals).

We analyzed our 32 core-set molecules, and in multiple independent experiments, found that 9 small molecules significantly extended the animals’ lifespan (in at least 3 to 4 trials, from ∼10% to ∼50%) (Supplemental Table 8). Gr-4D was the only non-toxic compound that consistently produced significant life-extending effects in 4 or more trials (average increase ∼24%; 50% in one trial) (Figure 4). Among the other 8 toxic molecules, O13, an orphan compound, also extended lifespan in a consistent manner (average increase ∼18%). Conversely, many molecules did not extend lifespan in any trial, and some even shortened lifespan, including two group-4 compounds (Gr-4A and Gr-4C) and O27 (Supplemental Table 8). In principle, lifespan might be increased due to caloric restriction; however, we did not observe an obvious reduction in pumping rates for worms treated with the lifespan-extending molecules (data not shown), and the animals were not pale, a hallmark of calorically-restricted animals.

To investigate the potential mechanism by which the chalcone Gr-4D extended lifespan, we asked whether it might affect gene activities known to influence longevity. To do this, we examined its effects on the expression of pathway-specific reporters, including the ER-stress reporter *Phsp-4::gfp*, the mitoUPR reporter *Phsp-6::gfp*, the NRF2/SKN-1 target gene reporter *Pgst-4::gfp* (for oxidative stress), and the autophagy reporter *LGG-1::GFP*. We observed that Gr-4D, at a dose that extended lifespan, caused modest induction of the *gst-4* transcriptional GFP reporter (Supplemental Figure 10), without exerting obvious effects on the others. As this glutathione transferase *gst-4* gene is up-regulated in response to SKN-1/NRF2 activation upon oxidative stress, these findings suggested that activation of SKN-1/NRF2 by Gr-4D might contribute, at least in part, to the lifespan extension in *C. elegans*. In the future, it will be interesting to perform genetic epistasis analysis of the life extension elicited by these compounds in *C. elegans*, as a way to learn more about underlying mechanisms.

## Discussion

To enrich for compounds that might slow aging, we screened for a cellular phenotype that is common to cells from many long-lived mutant animals and from naturally long-lived species of mammals and birds: resistance to oxidative stress. We screened ∼100,000 small molecules for their ability to protect a human primary fibroblast cell line from a lethal dose of hydrogen peroxide and retested the top hits for their ability to protect cells from the heavy metal cadmium and the DNA-damaging agent MMS. Many of these compounds conferred resistance to both hydrogen peroxide and cadmium. In addition, remarkably, ∼1/3 of our 32 core-set molecules extended *C. elegans’* lifespan (see the summary in Table 1).

Besides, 22 of the 54 repurchased compounds are not shown because: 1) Two orphan compounds (O19 and O25) scored negatively in all tests, even the hydrogen peroxide retest, and were discarded. 2) For three molecules (Gr-6A, O12 and O17), their masses did not match the predicted values by LC-MS analysis. 3) The other seventeen, including all group-2 compounds that were confirmed to be PARP inhibitors, increased ATP content upon stress (and potentially are still interesting), yet did not score positively for the cell death-imaging assay. Also, refer to Supplemental Table 1 for a list of the FOXO3 and NRF2 target genes assayed by qPCR.

### Properties of Hit Molecules

#### Chalcones

Four of our hits were chalcone-family members, including our most interesting hit G4-4D, which increased lifespan consistently in worms (up to ∼50%, 4 of 4 trials) with no apparent toxicity (Figure 4, Supplemental Table 8, also see Table 1).

Chalcones have been reported to produce many beneficial health effects, including anti-cancer, anti-HIV, anti-malarial, anti-inflammatory and anti-allergic activities (Batovska and Todorova, 2010; Sahu et al., 2012). They have a wide variety of molecular targets, many of which have relevance for cancer, which might explain their anti-proliferative activities against cell lines derived from many types of tumors (Solomon and Lee, 2012; Yadav et al., 2011).

Our 104K-compound screening library contained 71 chalcones (exact core-structure match), yet we only identified 4 among our 61 primary hits (group 4), suggesting an important role of side groups in determining H_2_O_2_-resistance (with the caveat that the others were not re-tested).

Interestingly, small molecules that contain an *α*,*β*-unsaturated ketone, the chalcone backbone, were identified recently in a screen for small molecules that promote proteome stability in mammalian cells (Calamini et al., 2011). Several proteins, including HSF-1, FOXO3 and NRF2, as well as the chaperone machinery, have been reported to mediate the beneficial effects of these small molecules. Of these, we only observed consistent signatures for NRF2-regulated gene, in both worms and human cells; however, genetic epistasis analysis indicated that, while NRF2 may potentially contribute to the stress resistance of human cells, NRF2-independent mechanisms play an important role as well.

Recently, Carmona-Gutierrez and co-workers identified 4,4’-dimethoxychalcone (DMC) as a natural compound that extended lifespan in multiple species, including yeast, worms and flies (Carmona-Gutierrez et al., 2019). The authors proposed that lifespan extension could be attributed DMC’s ability to induce autophagy. We did not observe obvious effects of our chalcones on autophagy in mammalian cells, using the Promega LC3 HiBiT assay system (with TOR inhibitor rapamycin as a positive control), nor did we observe significant effects of DMC in promoting resistance to H_2_O_2_ in WI-38 cells. The significance of this is not clear, as, for reasons we do not understand, we were also unable to observe an increase in autophagy with DMC in our assay system.

### Reported Properties of Hit Molecules

We referred to chemical databases (for example, PubChem) to cross-reference our small molecules to other high-throughput screens. We found that at least 9 molecules had been identified in multiple screens and suggested to affect certain human protein targets (Supplemental Table 9; See supplementary discussion). Of these, the chalcone Gr-4D, our top non-toxic compound, has been identified in many screens in the PubChem database. For example, it has been reported to kill fibrosarocoma cell lines that produce the oncometabolite 2-hydroxyglutarate (2HG) (AID 686970, with potency of ∼3 μM; see Supplemental Table 9). Other studies have suggested that Gr-4D can inhibit IL-1B-mediated inflammasome signaling (AID 743279) and activate p53 by inhibiting the MDM2/MDM4 interaction (in multiple assays). Finally, Gr-4D could induce DNA-rereplication in both SW480 colon adenocarcinoma cells and MCF 10A epithelial cells (with a potency of ∼5-6 μM), which in turn can induce the DNA damage response and apoptosis (BioAssay AID 624296 & 624297, reported target GMNN). Given the concern of genome instability that can be caused by DNA re-replication, it would be interesting to analyze whether a lower dose of Gr-4D could be tolerated *in vivo*.

Among the remaining 8 toxic compounds, Gr-3B and Gr-7C were identified in multiple assays that examined different biological targets (ranging from NK-*κ*B signaling, p53 regulation, potassium channel regulation, to protein SUMOylation). This raises a concern about non-specific actions of these molecules in cells. However, except for O6, which contains a reactive phenolic Mannich base, the other molecules do not appear to carry obvious PAINS signatures. By contrast, curcumin, a known PAIN molecule, has been found in at least 2167 assays with 92 reported protein targets (of these, 79 are human proteins, including NRF2, see Supplemental Table 9). Likewise, the other PAIN molecule resveratrol has been identified in at least 4266 assays with 91 active protein targets (these include 74 human proteins, such as SIRT1 and FOXO3).

To our surprise, despite the differences in their structures, several different molecules appeared to affect the same protein targets (Supplemental Table 9). For example, 4 molecules (the non-toxic chalcone Gr-4D and toxic Gr-1F, Gr-3B and Gr-7C) were shown previously to affect the cystic fibrosis transmembrane conductance regulator (CFTR), a chloride channel protein that is involved in multi-drug resistance (Borst et al., 1999). Like Gr-4D, 4 cell-toxic compounds (Gr-1F, Gr-3A, Gr-7C and O13) were also reported to target geminin GMNN, a DNA replication inhibitor. In addition, Gr-4D, Gr-3B and Gr-7C have together been identified in different studies and reported to inhibit multiple targets: for example, interleukin IL1B, a mediator of inflammasome signaling; MDM2 and MDM4, E3 ubiquitin protein ligases and negative-regulators of p53; NOD1 and NOD2, intracellular pattern-recognition receptors that regulate apoptosis and inflammation; and TDP1, a tyrosyl-DNA phosphodiesterase that interacts with PARP and regulates DNA damage response (see Supplemental Table 9). We do not know whether the ability of our molecules to promote oxidative-stress resistance is mediated, in part, by their effects on these reported target proteins.

#### Inhibition of NF-κB Signaling

Among the top non-toxic compounds, Gr-4D has been found in at least two screens linked to NF-*κ*B signaling. In one case, Gr-4D was reported to increase the expression of a luciferase reporter driven by the NF-*κ*B promoter in the SH-5YSY human neuroblastoma cell line (AID 1239). Curiously, it was also identified in another study as a molecule that inhibited TNF-alpha-induced NF-*κ*B activation in HEK 293T cells (AID 1852), leaving its regulation of NF-*κ*B in question. Among the remaining members of the 32 core-set compounds, Gr-3C was identified previously as a molecule that induced the NF-*κ*B inhibitor NFKBIA in two screens (EC_50_, 8.5 μM and 11.0 μM; see AID 317145 and 317143), consistent with the report from another screen suggesting it as an RELA inhibitor in HUVEC cells (IC_50_, 2.0 μM, AID 317146). These two studies above also identified Gr-3A, but not Gr-3B (which has a different side group that could introduce steric hindrance), as an NFKBIA-inducing compound (EC_50_, 11.0 μM and 12.0 μM). These findings are consistent with our analysis using the *in silico* docking database ZINC (Irwin et al., 2012), which predicted that Gr-3C bind to both NFKBIA and RELA (data not shown).

NF-kappaB signaling controls cell survival, differentiation, and proliferation. Key components of the pathway mediate inflammatory response (Lawrence, 2009) and have been implicated in many diseases, including cancer, autoimmune diseases, neurodegenerative diseases, cardiovascular diseases and diabetes (Hayden and Ghosh, 2012). Recently, it was shown that blocking age-related hypothalamic NF-*κ*B activation could retard aging and extend lifespan of mice (Zhang et al., 2013), thus potentially highlighting a role of chronic inflammation in aging (Jenny, 2012; Salminen et al., 2008). In fact, despite long-term cell toxicity observed for group-3 molecules, we noted that they all extended *C. elegans’* lifespan (3 of 3 trials, ∼8% to 33%), even though worms do not have apparent NF-*κ*B homologs.

### Hit Molecules as Regulators of Longevity

In animals, the rate of aging can be influenced by many factors, including a network of signaling proteins and transcription factors that also sense nutrients, energy levels and stress. Perturbing many genes in this network can extend healthspan and increase disease-resistance and lifespan (Bartke, 2011; Fontana et al., 2010). This is the case for long-lived *C. elegans* insulin/IGF-1 pathway *daf-2* mutants (Kenyon et al., 1993), where lifespan extension relies on DAF-16/FOXO’s regulation of a diverse collection of cell-protective, proteostasis, metabolic, innate-immunity and other genes (Kaletsky et al., 2016; Murphy et al., 2003; Schuster et al., 2010). This endocrine regulation of lifespan has been shown to be conserved among many different experimental species, and likely also small dogs and bats (Kenyon, 2010a).

FOXO is required for fly insulin/IGF-1 signaling mutants to live long (Slack et al., 2011) and for calorie restriction to extend mouse lifespan (Shimokawa et al., 2015). *AKT1* knockdown, our control that was expected to activate FOXO3, increased stress resistance in our assay. However, we identified only 3 molecules within our 32 core-set hits that may activate FOXO3, suggesting that our stress-resistance screen may not be ideal to look for FOXO3 activators.

Conversely, we observed an enrichment of small molecules that can activate NRF2. Increased activity of SKN-1/NRF2, the oxidative stress and xenobiotic phase II detoxification regulator, can extend life independently of *daf-16/foxo*. In addition, like *daf-16*, SKN-1/NRF2 promotes longevity in insulin/IGF1 pathway mutants (Tullet et al., 2008) and in calorically restricted worms (Bishop and Guarente, 2007), and also in fly *keap1* mutants (Sykiotis and Bohmann, 2008). It is activated in long-lived IGF1 pathway mouse mutants (Blackwell et al., 2015; Steinbaugh et al., 2012), and in long-lived mice lacking the glutathione S-transferase gene *mGsta4* (Singh et al., 2010). More recently, constitutive Nrf2 signaling activity has been found to be present in long-lived rodent species, including naked mole rats (Lewis et al., 2015), and enhanced cell signaling via Nrf2 and p53 have been suggested to be protective against spontaneous neoplasia and tumorigenesis in naked mole rats (Lewis et al., 2012).

We identified 14 potential NRF2 activators (3 non-toxic molecules, plus the non-toxic yet non-specific PAIN O6, as well as 9 other molecules with liabilities – see Results), of which at least 8 molecules (including 2 non-toxic, Gr-4D and Gr-5A; plus 6 other molecules with liabilities) increased resistance to both H_2_O_2_ and cadmium. 12 of the 14 molecules activated NRF2 without affecting FOXO3 or mTOR (Table 1); the two exceptions, Gr-7A & Gr-7C, were cell-toxic. Thus, NRF2 could be a significant contributor that promotes stress resistance in our screening conditions, consistent with the finding that NRF2 is activated in fibroblasts from long-lived animals, which also are resistant to multiple stressors (Leiser and Miller, 2010). That said, it was interesting that, while it might contribute to stress resistance, our epistasis analysis with *NRF2* siRNA knockdown indicated NRF2 was not required for the increased stress resistance of WI-38 cells produced by our small molecule hits.

The potential enrichment of NRF2-activating small molecules in our screen is also consistent with the anti-oxidative role of NRF2, and it may have implications for human aging as well. In human fibroblasts, reduced NRF2 function has been shown to occur in replicative senescence and *NRF2* silencing may lead to premature senescence (Kapeta et al., 2010). Conversely, Kapeta and co-workers also showed that molecules that activate NRF2 can enhance the survival of human fibroblasts following oxidative stress and extend their replicative lifespan. Interestingly, long-term exposure to the mTOR inhibitor rapamycin has been shown to increase mitochondria biogenesis and increase replicative lifespan of human fibroblasts, and these effects appeared to be mediated, at least in part, by p62/SQSTM1-associated degradation of KEAP1 and activation of NRF2 (Lerner et al., 2013). Similarly, multiple studies showed that rapamycin-induced lifespan extension requires SKN-1/NRF2 but not DAF-16/FOXO in *C. elegans* (Bjedov et al., 2010; Harrison et al., 2009; Robida-Stubbs et al., 2012; Wilkinson et al., 2012). Moreover, NRF2 activation also has been linked to several human age-related diseases, including atherosclerosis, neurodegenerative diseases, and certain types of cancer (Hybertson et al., 2011). Small-molecule modulators of the KEAP1-NRF2-ARE pathway are under development as potential preventive and therapeutic agents (Magesh et al., 2012), but whether their apparent ability to slow aging is compatible with disease resistance in humans remains to be seen.

### Hit Molecules as Potential Cancer Therapeutics

Many long-lived animal mutants are resistant to age-related diseases (Le Couteur et al., 2012), including cancer (Ikeno et al., 2009), protein-aggregation diseases (Cohen et al., 2009; Morley et al., 2002) and heart disease (Birse et al., 2010; Harrington et al., 2007; Wessells et al., 2004). Thus, the small molecules we seek may counter multiple age-related diseases.

Different species in nature have very different lifespans, but within each, the risk of cancer rises with age. This correlation in nature between slow aging and delayed cancer suggests that the same pathways that slow aging may antagonize cancer, and, in fact, many longevity pathways have anti-cancer activity (though some, such as Nrf2, can promote the growth of certain tumors). Importantly, these pathways inhibit a wide variety of cancers (Kalaany and Sabatini, 2009), which is what we predict for our small molecules. In fact, some of our stress-resistance hits, like the chalcones, are members of structural classes with known anti-tumor activity. When we compared the transcriptional signatures of cells treated with small molecules, we noted that several group-4 and group-5 molecules appeared to attenuate basal cell carcinoma signaling, based on the expression patterns of pathway genes consistent with prior knowledge (Supplemental Figure 7B). Interestingly, our non-toxic chalcone Gr-4D also appeared to attenuate signaling activity of the pro-inflammatory cytokine IL-8 (p value-adjusted, Figure 3B & Supplemental Table 6), which has been implicated in regulating a microenvironment favoring tumor progression.

Finally, even the small molecules that have cell toxicity could be interesting. In principle, any toxicity caused by these hits, which are administered at relatively high levels, could be off-target and thus dialed out via medicinal chemistry. Alternatively, they could be on-target toxicities. In the latter case, they could potentially increase stress resistance by activating cell-protective mechanisms, as do low levels of the herbicide paraquat in invertebrates, a situation (often called “hormesis”) that can extend lifespan in experimental organisms. Furthermore, toxicity could be cell type-specific. For example, we found that two group-7 molecules (Gr-7A & Gr-7B), which both produced long-term cell toxicity and appeared to inhibit mTOR, reduced the ATP content by ∼40-50% in a human lung adenosquamous carcinoma cell line HTB178 (data not shown). Conversely, when administered overnight at 10μM and assayed in parallel, these same molecules did not produce obvious effects on ATP levels in either the WI-38 primary cell line or in CRL-2081, lung-derived mesothelioma cells. Such selective effects could be due to differences in these tumor cell lines (for example, HTB178 cells carry mutations in EGFR and p53, while CRL-2081 cells have amplified MYC activity). These preliminary results also suggest an attractive possibility that some of our small molecules, when given at a dose that can be tolerated by normal cells, could selectively kill certain tumor cells. We should note that many cancer drugs, such as doxorubicin, can damage DNA and are intrinsically cell-toxic.

### Perspective

Our high-throughput screen in human primary cells was designed to enrich for stress resistance-promoting small molecules that could potentially perturb pathways that affect lifespan. We characterized the hit compounds in several different cellular assays, and *in vivo,* in *C. elegans*. By doing these assays, we found several molecules, most notably the chalcone Gr-4D, that could be of potential interest for further translational studies.

## Materials and Methods

### Small Molecule Screen Library

We screened a “diversity library” that contains 104,121 small molecules (∼24K from ChemBridge; ∼50K from ChemDiv; and ∼30K from SPECS) selected to maximize the coverage of chemical space. Compounds were provided as 10 mM stocks by the Small Molecule Discovery Center (SMDC), and chemical information, including structures, simplified molecular-input line-entry system (SMILES) IDs, PubChem links and possible *in silico* docking, is available for query. The SMILES of 209 primary screen hits are shown in Supplemental Table 1.

### Cell Culture & H_2_O_2_-resistance Screen

For the screen, we used human lung-derived WI-38 cells at mid population doubling levels (PDLs ∼30; these cells undergo replicative senescence at PDL ∼50). To reduce the variation due to difference in the doubling levels, cells were propagated from the initial stocks (ATCC, PDL ∼23) and prepared as frozen aliquots to be used for the screen as well as follow-up characterization. Prior to experiments, frozen cells were cultured and dissociated using Accutase (Sigma) for sub-cultivation at least once. In addition, human primary dermal fibroblasts (HDFp.05, from multiple donors) were obtained from Zen-Bio and cultured in dermal fibroblast medium, following the vendor’s instructions, and used for hit validation experiments. WI-38 cells were cultured in OptiMEM (Life Technologies) supplemented with 10% fetal bovine serum (FBS). In a 384 multi-well format, 2,000 cells were added in 50 μl medium per well, using a WellMate (Thermo Fisher) liquid dispenser and cultured in a CO_2_ incubator at 37°C for 24 hours. Then small molecules were introduced from stock plates with a 384-well formatted set of 50 nL stainless steel pin-tool (V&P Scientific) on a Biomek FXP (Beckman) automation workstation to 10 μM final concentration (in 0.1% DMSO), a typical single-concentration assay dose, which has been shown to produce high structural diversity in many small-molecule screens (Walters and Namchuk, 2003). The negative (0.1% DMSO-treated cells, n = 32) or positive controls (50 nM calyculin-treated cells, n = 32) were assigned to columns 1 & 2 or 23 & 24 respectively on the 384-well cell-culture plates to reduce potential cross-contamination with library compounds. Twenty-four hours later, the cells were subjected to a 3-hour stress treatment with 700 μM H_2_O_2_, and viability was assessed using CellTiter-Glo reagent (Promega), measuring luminescence signals on an Analyst HT (Molecule Devices) plate reader.

As our controls to knock down either *AKT1* or *KEAP1*, reverse siRNA transfection of WI-38 cells was performed, following an established protocol from the Schwarzbauer lab at Princeton with slight modifications. Briefly, siRNA oligos (Qiagen Flexiplate, validated) were complexed first with RNAiMAX (Life Technologies) and applied to 384-well microtiter plates, and then 2,000 WI-38 cells of mid-PDL were dispensed into the wells. Transfection was conducted first in medium with a lower serum level (∼6%), which was adjusted back to 10% final 4 hours later (50 μl, 20 nM siRNA final concentration), and then continued for ∼72 hours till the point for H_2_O_2_ stress and cell viability assays by measuring both ATP content and fraction of PI-positive cells. In the experiment to address *NRF2* dependency, WI-38 cells were transfected for ∼90 hours with double-stranded *NRF2* siRNA oligos (5’-UCCCGUUUGUAGAUGACAA-3’) (Singh et al., 2008) and then treated with Gr-4D for 24 hours before treatment with H_2_O_2_. Knockdown of *NRF2* was verified by RT-qPCR (normalized to the peptidylprolyl isomerase A gene *PPIA*).

To assess variation due to seeding, non-treated control plates were prepared and analyzed as well. The mean of standard deviations for all the control plates on multiple screen days was 10.8% ± 2.7% for non-stressed cells (data not shown). The average Z’ value, measuring the difference between positive and negative controls to assess the extent of variation, was 0.61 (Supplemental Figure 2), a robust value for such a screen. We discarded molecules that significantly increased the ATP signal by stimulating cell proliferation in the absence of H_2_O_2_.

To perform the original dose-response and cell-death imaging analysis, 209 candidate hits were picked individually from the screen library and re-analyzed at six different final concentrations (0.6 μM, 1.25 μM, 2.5 μM, 5 μM, 10 μM and 20 μM) to examine their ability to promote H_2_O_2_-resistance of WI-38 cells. Cells pre-treated with these molecules (at 1.25 μM and 10 μM) were also analyzed by propidium-iodide imaging to examine cell death following 3 hours of H_2_O_2_ treatment. For DMSO pre-treated controls (n = 30), the percentages of PI-positive cells were 23.7% ± 6.9% (average ± standard deviation, 1.25 μM assay plate) and 27.4% ± 6.5% (average ± standard deviation, 10 μM assay plate). When assayed at 1.25 μM and 10 μM, 107 hits were found to reduce the percentage of PI-positive dead cells (by 1 to 3 standard deviations from the mean) upon H_2_O_2_ (data not shown). Note that certain effective small molecules could have been missed in the imaging assays, as false negatives could arise due to potential stability issues of the library compounds. Fresh small molecules were obtained from different vendors, including ChemBridge, ChemDiv, Vitasreen and MolPort, and analyzed by LC-MS for quality validation. Except for three molecules (Gr-6A, O12 and O17), molecular masses were confirmed to match with predicted values.

To determine whether hit compounds possess any structural similarity to each other and/or to chemical moieties with known properties, we used the MetaDrug chemical data-mining tool (Ekins et al., 2007) and the Similarity Ensemble Approach (SEA) statistical method (Keiser et al., 2007), which compares compounds to known therapeutic drugs.

### CdCl_2_ resistance Assays

Small molecules were tested for cadmium resistance twice in two independent assays. Each small molecule was analyzed at five different concentrations (0.25 μM to 20 μM final concentration), with technical triplicates for each dose. Data were analyzed using a global variance t-test. The experimental setups on 384-well plates and small molecule incubation times were the same as for H_2_O_2_ assay, and the incubation time was 12 hours for both cadmium (700 μM) and MMS (900 μM). We tested several conditions, including conditions used to assay fibroblasts from long-lived animals (Salmon et al., 2005). These conditions produced better results with the positive controls (*AKT1* and *KEAP1* siRNAs) in the MMS assay. Thus, for MMS, the cells were shifted from growth media – OptiMEM plus 10% FBS, to DMEM (Gibco) plus 2% BSA (no serum) just before adding small molecules, which occurred 24 hours before MMS addition.

### DPPH Assay

2,2-diphenyl-1-picrylhydrazyl (DPPH) is a radical-containing purple dye that can be reduced by ROS scavengers. This cell-free assay was performed as described previously (Sharma and Bhat, 2009). DPPH (Sigma) stock (25 mM) was prepared in methanol and diluted to 50 μM in acetic acid-buffered methanol (0.1 M, pH 5.5). 50 μl diluted solution was dispensed into three 384-well assay plates (technical triplicates). 209 small molecule hits were picked individually from the library stock onto a stock plate, and then introduced by a pin-tool into the assay plates at 10 μM final concentration. DMSO negative control and several positive control ROS scavengers, including N-acetyl cysteine, amodiaquin dihydrochloride and 8-hydroxyquinoline quinoline (8-HQ) (all from Sigma), were introduced into separate wells. Plates were sealed and incubated in a humid chamber at 30° C in the dark. Absorbance at 519 nm was measured on a FlexStation 3 multi-mode microplate reader (Molecular Devices) 3 hours and 24 hours later.

### AmplexRed Assay

This cell-free assay was performed using the Amplex Red Hydrogen Peroxide/Peroxidase Assay Kit (Life Technologies), following the manufacturer’s instructions. Briefly, small molecules (at 10 μM final concentration) were pre-incubated with 700 μM H_2_O_2_ in water for 3 hours in a 37°C CO_2_ incubator. Catalase (MP Biomedicals) was used as the positive control. Amplex Red reagent was mixed with horseradish peroxidase (HRP) in buffer and then incubated with 1:10 diluted small molecule/ H_2_O_2_ mixture for 30 minutes in dark. Fluorescence at 590 nm was measured on a FlexStation 3 multi-mode microplate reader (Molecular Devices).

### Imaging Analysis

Cells were pre-incubated with small molecules for 24 hours and then subjected to H_2_O_2_ for 3 hours (in several experiments, multiple, consecutive times points were included for analysis). Hoechst 33342 (Life Technologies) (10 μg/ml final concentration) and propidium iodide (Life Technologies) (2.5 μM final concentration) were added for the last 30 minutes. Fluorescent images of cells were collected on the INCell Analyzer 2000 automated microscope (GE Healthcare) (10X objective) and further analyzed with the Developer ToolBox software (GE Healthcare; version 1.9).

### Cell Confluency Analysis

WI-38 cells (2,000 per well) were seeded on 96-well plates and cultured and scanned every 2 hours to record their confluency (relative percentage of surface area in a cell-culture vessel covered by cells) in an IncuCyte Zoom Live-Cell Analysis System (Essen Bioscience) for up to 112 hours. Small molecules (10 μM final concentration, n = 3 for each small molecule) were introduced at 24 hours following the start point. Relative cell confluency (by surface area of a given well) was analyzed with the vendor’s software.

### DNA Damage Marker Analysis

WI-38 cells were seeded (∼8,000 cells per well) on 96-well plates, cultured for 24 hours and then incubated with small molecules for another 24 hours. Doxorubicin (300 nM, 24 hours) or H_2_O_2_ (700 μM, 3 hours) was also individually introduced as the positive control to damage DNA. Cells were washed with phosphate-buffered saline (PBS), fixed with 4% paraformaldehyde for 30 minutes and then blocked with 5% normal goat serum (Cell Signaling) and 0.3% Triton X-100 for 1 hour. Primary antibody cocktail [Cell Signaling, rabbit anti-phospho-histone *γ*H2A.X (Ser-139), 1:100; anti-phospho-TP53BP1 (Ser-1778), 1:100] was prepared in PBS with 1% BSA and 0.3% Triton X-100 and incubated overnight at 4°C. The next day, samples were washed with PBS and further incubated with fluorophore-conjugated secondary antibody (1:1,000) for 1 hour in dark. Samples were then washed and incubated with DAPI dye (Life Technologies) (4 μg/ml final concentration) for 30 minutes. Images were collected on the INCell Analyzer 2000 (10X objective) and analyzed with the Developer Toolbox. Cells that showed immuno-staining intensity above a software-defined threshold were scored.

### PARP Inhibitor Assay

The assay was performed with the HT Fluorescent Homogeneous PARP Inhibition Assay Kit (Trevigen), following the manufacturer’s instructions. Briefly, nicotinamide adenine dinucleotide (NAD), human PARP1 and activated-DNA solution were distributed across a 96-well plate. Fifty-one repurchased, mass-checked small molecules were introduced at 10 μM final concentration and incubated in a humid chamber at room temperature for 30 minutes in the dark. Cycling mixture, with resazurin and cycling enzyme diaphorase, was then added and incubated further for 1 hour in dark. The reaction was terminated with stopping buffer. Fluorescence was then measured on a FlexStation 3 multi-mode microplate reader (544 nm excitation/ 590 nm emission).

### RNA-seq Analysis

For RNA-seq analysis, WI-38 cells were treated with Gr-4D (2.5 μM final, a dose confirmed to promote H_2_O_2_-resistance in a paralleled experiment – – see Supplemental Figure 6) from two different vendors (CB, ChemBridge; MP, MolPort), 0.1% DMSO control or no DMSO mock control (n = 8 each) in the absence of H_2_O_2_ for 24 hours and then processed for RNA isolation with the RNeasy kit (Qiagen). Total RNA (375ng, normalized) was used for each sample to prepare Illumina TruSeq Stranded mRNA library, following the manufacturer’s instructions. Multiplexed samples were analyzed on an Illumina HiSeq 4000 to obtain transcript reads. The resulting reads were processed using our in-house analysis pipeline to obtain transcript abundance. Genes that showed significant change of expression (normalized to DMSO-treated control samples) were identified (with false discovery rate < 0.05, 1.5-fold) and used for pathway analysis, using tools including PANTHER (v14.0, http://pantherdb.org/) and Enrichr (http://amp.pharm.mssm.edu/Enrichr/). The iLINCS database (http://www.ilincs.org/ilincs/signatures/main/upload) was used for comparison analysis to identify perturbations that produced expression profiles similar to our small molecules.

### Microarray Analysis

For microarray analysis, WI-38 cells were treated with small molecules (10 μM final, a dose confirmed to promote H_2_O_2_-resistance in a paralleled experiment – see Supplemental Figure 6; n = 3 each for group-4 and group-5 molecules, plus O10) or 0.1% DMSO control (n = 15) in the absence of H_2_O_2_ for 24 hours and then processed for RNA isolation with the RNeasy kit (Qiagen). Reverse transcription (RT) and production of Cy5-labeled cRNA and further hybridizations were performed with the Agilent two-color microarray kit (SurePrint G3 human v3 arrays, 8×60K), following the manufacturer’s instructions. Expression levels of gene probes were obtained by normalizing their absolute signals, using the Cy3-labeled Universal Human Reference cRNA (Agilent). Genes that showed significant change of expression (normalized to DMSO control-treated samples) were obtained using Statistical Analysis of Microarray (SAM, with false discovery rate < 0.10, 1.5-fold) with the Multi Experiment Viewer (version 4.8). Normalized expression levels and FDR values for significant genes were further used for pathway analysis. Gene clustering analysis for microarray data was performed using Gene Cluster (version 3.0).

### qPCR Analysis

For qPCR analysis, WI-38 cells were treated with small molecules (10 μM final concentration, n = 4) for 24 hours and processed for RNA isolation and reverse transcription with the Cells-to-Ct Kit (Life Technologies), following the manufacturer’s instructions. RT products were diluted with H_2_O and used for qPCR analysis on an ABI 7300 system (Life Technologies) (technical triplicates). Relative expression levels of target genes were calculated by the ΔΔCt method, using the reference gene beta-2-microglobulin *B2M*, and relative fold changes were obtained by normalizing to negative controls and further analyzed using the Student’s t-test. We analyzed four NRF2-regulated genes, including *HMOX1* (heme oxygenase (decycling) 1; an anti-oxidant), *NQO1* [NAD(P)H dehydrogenase, quinone 1; a phase 2 detoxification enzyme], *GCLC* (glutamate–cysteine ligase, catalytic; a glutathione-synthesis enzyme) and *GSTM1* (a glutathione S transferase). We also analyzed another five FOXO3A-regulated genes: *SOD2* (superoxide dismutase), *GADD45A* (a cell cycle regulator), *CAT* (catalase), *DDB1* (damage-specific DNA binding protein) and *TXNIP* (thioredoxin-interacting protein). Sestrin 1 (*SESN1*), a gene known to be regulated by both NRF2 and FOXO3A, was also analyzed by qPCR.

### mTOR Inhibition (RPS6 Phosphorylation Status) Analysis

In-Cell Western assays were performed, following a standard immuno-staining protocol. Briefly, WI-38 cells were treated with small molecules for 24 hours, and then processed and incubated with primary antibody cocktail (Cell Signaling, mouse anti-RPS, 1:25; rabbit-anti-pRPS6-Ser-235/236, 1:100) overnight at 4° C. The next day, samples were processed and incubated with fluorophore-conjugated secondary antibodies (Cell Signaling, DyLight 680-goat anti-mouse, 1:500; DyLight 800-goat anti-rabbit, 1:1,000). Images were collected on an Odyssey Imager (LI-COR) and analyzed with the Image Studio Lite software (version 5.0.21).

### Poly(Q) Toxicity & Viability Analysis

Assessment of poly(Q) toxicity was performed as described previously (Aiken et al., 2004). PC12 cells that stably express the inducible poly(Q)103-Htt-EGFP were grown in culture for 24 hours and then subjected to the treatment with small molecules (10 μM). Ponasterone A (Life Technologies), an ecdysone analog, was introduced at 10 μM 24 hours later to induce transgene expression, and the formation of aggregation puncta was examined and confirmed using the Eclipse 200 fluorescent microscope (Nikon). Cell viability was analyzed 48 hours later following the induction, by measuring ATP content with CellTiter-Glo. The parental WT-PC12A cells, which do not express poly(Q)103-Htt-EGFP, were used as the negative control to exclude the possibility that certain small molecules may enhance ATP content even in the absence of toxic aggregates.

### Lifespan Assays in C. elegans

Given the caveat of lifespan-assay variations for *C. elegans* studies, lifespan analysis was carried out using several different culture conditions (in liquid and on plate, food concentrations, live or UV-& kanamycin-treated bacteria). Molecules were analyzed at the highest dose (∼60 μM, 0.3% DMSO, as higher DMSO concentration has been reported to extend lifespan of *C. elegans*). Liquid culture-based lifespan assays were performed, following the protocol as described (Solis and Petrascheck, 2011). Briefly, synchronized L1s of wild-type animals were fed ampicillin-resistant OP50 bacteria and treated with small molecules (∼60 μM final concentration, 0.3% DMSO) at the young-adult stage. FuDR was used at 100 μM final concentration at the L4 stage to block progeny production. The molecules were analyzed in 96-well plates, with 4 wells for each small molecule. Multiple control wells with DMSO (0.3% final concentration) were included on each plate. Likewise, small molecules were also analyzed for their ability to extend lifespan on solid agar, following procedures as described (Cabreiro et al., 2013). Hypochlorite-synchronized temperature-sensitive sterile mutants, CF4059, [*fer-15(b26)II rol-6(su1006)II; fem-1(hc17)IV*], were raised on large agar plates seeded with OP50 bacteria at 25°C. Sterile day-1 adults were transferred onto mini-plates, which were seeded with normal OP50 bacteria (UV-irradiated, kanamycin-treated) and supplemented with small molecules (∼60 μM final concentration, 0.2% DMSO). Worms were scored every other day. Cumulative survival was analyzed using the STATA software (log-rank test).

### H_2_O_2_ Stress Assays in C. elegans

This stress assay was performed in liquid, following procedures similar to lifespan assays as described above. Wild-type worms were treated with the small molecules from young adulthood, and then H_2_O_2_ (0.5 mM final concentration) was added on day 8 (1^st^ trial to test all the molecules repurchased) or day 6 (2^nd^ trial to retest candidate hits from the 1^st^ trial) of adulthood, and then animals were scored for viability every day.

### Effects on pathway activity reporters in C. elegans

Transgenic worms expressing different reporters of several longevity-related pathways were raised from the L4 stage on mini-plates that were seeded with normal OP50 bacteria, supplemented with 20uM FuDR (to block progeny production, and supplied with either DMSO (as the control) or different compounds. About 30 hours later, at least 12-15 adult worms per condition were imaged on a Nikon Eclipse Ti spinning disk confocal microscope (10x objective, 408nm).

## Data availability

Supplemental materials (tables and figures) are available at *Figshare*. Table S1 shows the chemical identities of 209 primary hits, plus 61 top hits and 32 core-set hits analyzed. Table S2 shows the cell death-imaging analysis to confirm protective effects of small molecules against H_2_O_2_. Table S3 shows that certain small molecules also protect cells from heavy metal cadmium. Table S4 indicates the effects of small molecules on DNA damage-associated markers and ATP contents upon prolonged incubation. Table S5 lists the significant genes identified by RNA-seq (for Gr-4D) and microarrays (for other molecules). Table S6 shows the pathway analysis results. Table S7 shows that certain molecule might be able to inhibit mTOR. Table S8 summarizes the effects of small molecules on *C. elegans’* lifespan. Table S9 lists potential targets for our small molecules.

Figure S1 shows increased oxidative-stress resistance upon *AKT1* or *KEAP1* knockdown. Figure S2 shows Z-prime scores across the screen. Figure S3 indicates the analysis to exclude ROS scavengers. Figure S4 shows no obvious quenching of H_2_O_2_ by small molecules *in vitro*. Figure S5 indicates the long-term effects of small molecules on ATP levels and cell confluency. Figure S6 shows increased cell viability upon H_2_O_2_ by small molecules in the RNA-seq and microarray experiment, as well as *NRF2* non-dependency for Gr-4D. Figure S7 displays the effects of small molecules on gene expression of WI-38 cells. Figure S8 indicates the analysis to identify certain PARP inhibitors. Figure S9 shows protective effects of certain small molecules against poly-Q toxicity. Figure S10 shows the effects of Gr-4D on *C. elegans* expressing reporters of genes related to pathways known to influence longevity.

## Acknowledgements

PZ and JC conducted the screen, with crucial advice and assistance from KA and MA. PZ, YZ and JC characterized the screen hits. PZ and CK designed the experiments and prepared the manuscript. We thank SMDC staffs for their assistance to analyze the repurchased small molecules by LC-MS. We are grateful to Erik Schweitzer (UCLA) for the PC12 stable cell lines expressing poly(Q)-HTT::EGFP and Michael Petrascheck (Scripps) for the ampicillin-resistant OP50 bacteria. The MK-4827 PARP1 inhibitor was a kind gift from Xiaodong Yang and Daphne Haas-Kogan (UCSF). We also thank Katie Podshivalova for generous help with data analysis and critical reading of the manuscript. Finally, we thank the Genomics Core and data platform team at Calico for their kind help with RNA-seq analysis. This work was supported by philanthropic funding from the Thiel Foundation and a NIH/NIA grant (R01 AG044515) to CK. All 61 hits that passed the initial selection and their characterizations have been disclosed to UCSF (see Patent US20180289682A1).

## Supplemental Text

### Addressing Long-term Effects of Small Molecules

We first performed viability assays to address the long-term effects of the molecules on WI-38 cells, by measuring ATP levels and scoring cell death during a course of 5 days of continuous treatment (in the absence of H_2_O_2_). Of 32 core-set molecules, we found at least 11 that, compared with controls, reduced ATP content by more than 30% by day 5 of treatment (Supplemental Figure 5-1), suggesting potential anti-proliferative activity and/or cell toxicity of these molecules. Consistent with this, as indicated by the examination of cell confluency, morphology and cell death-imaging, 9 of these 11 molecules were found to reduce cell number and increase the fraction of PI-positive dead cells, and most cells treated with the remaining two (Gr-7A & Gr-9B) were dead (data not shown). Conversely, rapamycin (5 μM), a potent inhibitor of mTOR that reduces growth rates, reduced the ATP content and cell confluency by ∼50%, yet did not significantly increase cell death.

We also performed cell confluency-based proliferation assays, focusing on the small molecules that did not show obvious toxicity in the experiment described above. We found that of 14 small molecules analyzed, at least 5 did not show strong inhibitory effects on cell proliferation (Supplemental Figure 5-2). Consistent with the previous analysis, the other 9 small molecules that inhibited cell proliferation, unlike rapamycin, produced cell toxicity (assessed by cell morphology). We noted that all group-7 molecules and multiple members of group 4, except for one (Gr-4D), were toxic, suggesting that the same attributes that elicit stress resistance may cause toxicity (“on target” effects).

### Analyzing Potential DNA-damaging Effects of Small Molecules

One (undesirable) way in which a small molecule could induce stress resistance is by blocking apoptosis in response to DNA damage. Therefore, we examined effects on two DNA damage-associated cellular markers, phosphorylated histone variant *γ*H2A.X and tumor protein p53 binding protein 1 (TP53BP1) in WI-38 cells. *γ*H2A.X is required for checkpoint-mediated cell cycle arrest and DNA repair following double-stranded DNA breaks, and phosphorylation of *γ*H2A.X by a group of PI3K-like kinases (ATM, ATR, and DNA-PK) occurs rapidly in response to DNA damage (Perez-Cadahia et al., 2010). Likewise, in response to DNA damage, TP53BP1 is phosphorylated and translocated into the nucleus, and retention of TP53BP1 at DNA breaks requires phosphorylated *γ*H2A.X (Panier and Boulton, 2014). Among 32 core-set hits, seven small molecules (three members of group 4, two members of group 5, and two orphans) did not appear to increase the fraction of cells scoring positively for either marker in the absence of H_2_O_2_ stress (Supplemental Table 4). Consistent with this observation, 5 molecules (Gr-4D, Gr-5A, Gr-5D, O10 and O14) did not show strong cell toxicity in the cell-proliferation assays (Supplemental Figure 5). Conversely, the remaining molecules increased the fraction of cells scoring positively for at least one marker in one experiment (Supplemental Table 4).

We also note that, of the small molecules that increased *γ*H2A.X and/or TP53BP1 foci, 13 molecules also increased the percentage of propidium iodide-positive death cells under normal conditions (Supplemental Table 2), suggesting that cell toxicity is due to their effects to increase DNA damage and/or DNA damage-associated markers. In other words, a number of small molecules identified in our screen, likely by inducing modest levels of cellular stress (some, by increasing DNA damage), may protect cells from H_2_O_2_ through a “hormesis” mechanism, or, alternatively by preventing the apoptosis that would normally occur in response to DNA damage. Increased DNA damage can significantly elevate the risk of malignant transformation when affected cells do not undergo senescence and apoptosis. However, we noted that these DNA-damaging small molecules could still be interesting, as they may act like certain cytotoxic agents (e.g., doxorubicin) and kill highly proliferative tumor cells *in vivo*.

### “PAINS”

One general concern about small-molecule screens is whether the hits could have non-specific effects in cells. In particular, reactive electrophilic “pan assay interference compounds” (PAINS), can have highly pleiotropic effects on cells (Baell and Walters, 2014; Baell and Holloway, 2010). A typical academic screening library could have ∼5-12% of compounds that are PAINS, including catechols, rhodanines, phenolic Mannich bases, enones and others, which could behave as redox cyclers, metal complexers and covalent modifiers. Notably, different classes of PAINS have been observed to co-segregate frequently in the same biological screens. When we queried the diversity chemical library that we screened, using the known signature structures of PAINS, we found ∼18,191 possible PAINS, representing 17.5% of 104,121 total molecules. We further checked our small molecules and found that 7 of 61 primary hits (including a PARP inhibitor) and 5 of 32 core-set hits, contained at least part of the PAINS moieties (Supplemental Table 1). Among our top non-toxic hits, one, O6, was a PAIN (phenolic Mannich base); and hence, we excluded this molecule from microarray analysis. In addition, four of the toxic core-set hits exhibited PAINS signatures. Of these, Gr-6C contains a rhodanine moiety and Gr-9B shows both enone and catechol structures, and they extended *C. elegans*’ lifespan in multiple trials (Supplemental Table 8). Like O6, Gr-9A (enone) induced both *HMOX1* (by almost 50 folds) and *NQO1*, two NRF2-regulated genes that are involved in the xenobiotic stress response. As mentioned earlier, Gr-9A also protected worms from hydrogen peroxide. O21, another possible PAIN molecule with a rhodanine structure like Gr-6C, scored positive for DNA-damage markers (Supplemental Table 4).

Nonetheless, our screen did not seem to enrich for PAINS (*P* = 0.068, hypergeometric distribution probability, for all 61 hits). In the case of the potential PAINS, they produced many interesting phenotypes, including enhancing stress resistance, inducing expression of NRF2- and FOXO3-regulated genes, and extending lifespan of *C. elegans*. We also note that curcumin and resveratrol, two prominent PAINS (see Supplemental Table 9), extend lifespan in several species (Alavez et al., 2011; Hubbard and Sinclair, 2014) and also have potential value for treating cancer, diabetes and other diseases in humans (Anand et al., 2008; Brasnyo et al., 2011). In this regard, the PAIN molecules may still produce global benefits in a specific biological or pathological context in the whole animal.

### PARP Inhibitors

Poly ADP-ribose polymerase mediates cellular responses to many types of stress, including oxidative, genotoxic, inflammatory and metabolic stress (Luo and Kraus, 2012). PARP inhibition has been reported to increase cells’ resistance to DNA-damaging agents like hydrogen peroxide (Zhang et al., 2007). We recovered two compounds (group 8) that are highly similar to the PARP inhibitor 4-amino-1,8-naphthalimide (Banasik et al., 1992), an experimental cancer drug (Costantino et al., 2001). This prompted us to analyze all of our 51 repurchased and validated molecules in an *in vitro* PARP assay. In addition to these two group-8 molecules, 8 small molecules (including 4 members of the structural group 2) also substantially inhibit human PARP1 activity (Supplemental Figure 8). Of these, Gr-2E is a non-specific PAIN molecule. However, we noted that, despite their ability to preserve ATP content, only 4 of these molecules modestly reduced the percentage of PI-positive cells upon H_2_O_2_ stress (Supplemental Table 2). Thus, PARP inhibitors from our screen did not behave as agents that protect cells from oxidative stress, which is why we excluded these molecules from most of our analyses.

On the other hand, several recent studies suggest that PARP inhibition may promote health and longevity. First, deletion of *Parp1* has been reported to increase mitochondrial metabolism through NAD^+^ preservation and SIRT1 activation, and to protect animals from metabolic disease (Bai et al., 2011). Second, long-term treatment with PARP inhibitors has been found to enhance mitochondrial function and improve skeletal-muscle in mice, and also reverse mitochondrial defects in primary myotubes of obese humans and attenuate metabolism defects in *NDUFS1* (a mitochondrial NADH dehydrogenase and oxidoreductase that transfers electrons from NADH to the respiratory chain) mutant fibroblasts (Pirinen et al., 2014). Third, disruption of the *C. elegans* ortholog *pme-1/parp-1*, as well as treatment with PARP inhibitors, induced the mitochondrial unfolded-protein response (mtUPR) and extended lifespan in a manner that required both *sir-2.1* and *daf-16/foxo* in *C. elegans* (Mouchiroud et al., 2013). Consistent with the previous reports, we also found that some of our PARP inhibitors extended *C. elegans’* lifespan (specifically, Gr-8A, ∼10% to 34%, 4 of 4 trials; O14, ∼6% to 14%, 3 of 3 trials) (Supplemental Table 1).

PARP inhibitors can kill certain types of tumors because of their effects on DNA repair pathways (Chan et al., 2010; Mason et al., 2014; Rouleau et al., 2010); specifically, they have been shown to be effective in clinical trials among cancer patients carrying *BRCA1/2* mutations (Fong et al., 2009). The PARP inhibitors we identified could be interesting in this regard.

### Previously Reported Properties of Hit Molecules – *Inhibition of SUMOylation*

Among our 32 core-set hits, two molecules (Gr-1F & O13, both toxic) have been identified in several screens and confirmed to inhibit multiple SUMO/sentrin-specific peptidases (SENPs) (∼50-90% inhibition at 5-20 μM). Small ubiquitin-related modifiers (SUMOs) are ubiquitin-like proteins that can be attached covalently to a variety of target proteins (Geiss-Friedlander and Melchior, 2007). SUMOylation modification can by removed by SENPs, a class of SUMO-specific proteases, which affect many cellular processes, including apoptosis, DNA damage repair, ribosome maturation, and transcription (Hickey et al., 2012; Yeh, 2009). SUMOylation of many proteins increases remarkably in response to heat shock and hydrogen peroxide (Zhou et al., 2004), which may explain, at least in part, the mechanism by which our molecules promote stress resistance. Our discovery of potential SENP inhibitors could be interesting, as these molecules have the potential in treating certain diseases including cancer, atherosclerosis and heart diseases, where perturbed SUMOylation balance has been observed (Kumar and Zhang, 2015).

## Supplemental Figure Legends

Supplemental Figure 1 (related to Figure 1). Increased oxidative-stress resistance upon *AKT1* or *KEAP1* knockdown.

A) To define the optimal assay conditions for our screen, WI-38 cells were incubated with different doses of H_2_O_2_ for different periods of time in 384-well plates and then assayed for viability by measuring the ATP content. Shown here are representative data, indicating that high doses (above 600 μM) of H_2_O_2_ substantially depleted the ATP content after 3 hours of incubation (luminescence values are shown by the side). Control, n = 16; for all other H_2_O_2_ test samples, n = 11-16 (***, *P* <0.0001 for 500 μM H_2_O_2_ and above, Student’s t-test). Shown in all the figures, unless otherwise specified, were mean ± SD.

B) WI-38 cells were transfected with 20 nM AllStars negative control or *AKT1* or *KEAP1* siRNA oligos (Qiagen) and then assayed for viability 72 hours post-transfection upon H_2_O_2_. Representative data indicated that *AKT1* or *KEAP1* knockdown significantly increased cell viability, as measured by ATP content following 3 hours incubation in 700 μM H_2_O_2_. One-way ANOVA was performed for statistics (p<0.001), plus post-comparison with control (***, p<0.001). Control, n = 24; *AKT1* knockdown, n = 18; *KEAP1* knockdown, n = 13. Viability was also measured in the same experiment for cells not subjected to H_2_O_2_. One-way ANOVA (p<0.001), plus post-comparison with control (**, p<0.01; n.s., not significant). Control, n = 16; *AKT1* knockdown, n = 12; *KEAP1* knockdown, n = 12.

C) Propidium iodide staining was performed in parallel to analyze the fraction of dead/dying cells upon H_2_O_2_. Consistent with the ATP assay results, *AKT1* or *KEAP1* knockdown significantly reduced the fraction of PI-positive cells. One-way ANOVA (p<0.001), plus post-comparison with control (***, p<0.001). Control, n = 8; *AKT1* knockdown, n = 6; *KEAP1* knockdown, n = 6.

Supplemental Figure 2 (related to Figure 1). Z’ scores for the ATP assay across the screen. Shown is the Z’ score for each of the 327 plates carrying a total of 104,121 library compounds screened. Z’ score, defined as 1 – [(3X standard deviation for positive controls + 3X standard deviation for negative controls)/(mean value for positive controls - mean value for negative controls)], is typically used to access the assay quality in a high-throughput screen, and assay robustness is indicated by a Z’ score greater than 0.5. The positive control calyculin (EMD Biosciences), a potent serine/threonine protein phosphatase inhibitor, significantly increased the levels of ATP in H_2_O_2_-stressed WI-38 cells, relative to the DMSO negative control. Average Z’ score is 0.61±0.13 (mean ± SD) in our primary screen.

Supplemental Figure 3 (related to Figure 1). Cell-free ROS-scavenging assay of known ROS scavengers and our small molecules. The screen hits were analyzed together with several known ROS scavengers, including N-acetyl cysteine (NAC), amodiaquin dihydrochloride (AmD) and 8-hydroxyquinoline quinoline (8-HQ), in the absence of cells. Of 209 screen-hits assayed (at 10 μM), 56 molecules (including 40 that share a core structure of 8-HQ) were found to reduce the absorbance by 10% or more 24 hour-post incubation and were classified as putative ROS scavengers. The majority of these reduced the absorbance by ∼30%. Across 3 assay plates: 0.4% DMSO negative control, n = 417; each positive control of each dose, n = 12; each library compound, n = 3. Mean of absorbance normalized to DMSO control for each group was shown above.

Supplemental Figure 4 (related to Figure 2). No *in vitro* H_2_O_2_-quenching effects by small molecules. Catalase (0.02 units and 0.04 units), the positive control, substantially reduced the fluorescence in the Amplex Red assay that measures H_2_O_2_ concentration. By contrast, none of the 51 repurchased molecules appeared to reduce the absorbance. Across 4 assay plates: 0.2% DMSO negative control, n = 8; catalase positive control or each repurchased small molecule, n = 4.

Supplemental Figure 5 (related to Figure 3). 5-1) Long-term effects of small molecules on ATP levels of cultured WI-38 cells. Of 32 core-set hits, 28 molecules were analyzed in multiple 384-well plates in replicates in two batches, (A) 26 and (B) 2, to assess their effects on ATP levels upon prolonged incubation in the absence of H_2_O_2_ for up to 5 days (10 μM, n = 6 for each molecule, average standard deviation across the whole assay is ∼5.9%). Four molecules (Gr-3A, Gr-4B, Gr-4D and O6) were not included in this experiment, since likely due to compromised stability, these 4 of this specific batch did not retest for H_2_O_2_-resistance on day 2 of treatment. However, Gr-4B and Gr-4D from a new batch purchased were analyzed in the IncuCyte analysis of cell confluency (see Supplemental Figure 5-2). Green mark on the y-axis indicates the start-point ATP level measured for 1,000 cells 24 hours post-seeding, before adding any small molecules. Note that compared with 0.1% DMSO control (black thick line) and H_2_O control (aqua thick line), 5 μM rapamycin (yellow thick line), which is known to reduce cell proliferation and cell size, reduced ATP level by ∼50% on day 5 of treatment. Dashed lines: of 28 analyzed, at least 11 small molecules (Gr-1F, Gr-3B, Gr-3C, Gr-6C, Gr-7A, Gr-7B, Gr-7C, Gr-9B, O11, O21 and O27) reduced the ATP level by more than 30% by day 5 (see Supplemental Table 4 for details). These molecules also were examined in the same experiment for their effects on cell morphology and cell death by PI-imaging on day 2 and day 5 of treatment. Unlike rapamycin, which also reduced ATP levels substantially, 9 of these 11 molecules (Gr-3B, Gr-6C, Gr-7A, Gr-7B, Gr-7C, Gr-9B, O11, O21 & O27) also produced cell toxicity (by the examination of cell morphology, data not shown). The other two, Gr-1F and Gr-3C, caused DNA damages, like the others (see Supplemental Table 4).

5-2) Effects of prolonged small-molecule incubation on WI-38 cell confluency. Shown is a representative plot of cell confluency for each treatment condition. Confluency of WI-38 cells was monitored in an IncuCyte Zoom Live-Cell Analysis System for 112 hours. Small molecules (10 μM final, n = 3 each) were introduced at 24 hours (indicated by arrows) following the start point. Fourteen different small molecules that did not produce obvious long-term cell toxicity (see Supplemental Figure 5-1) were retested (vendors are listed at the bottom). A) As a positive control, the TOR inhibitor rapamycin (2.5 μM) reduced cell confluency significantly. B) 5 molecules did not show strong inhibitory effects on cell confluency, and C) 9 molecules significantly reduced cell confluency.

Supplemental Figure 6 (related to Figure 3). Increased cell viability upon H_2_O_2_ by small molecules in the RNA-seq and microarray experiment, as well as *NRF2* non-dependency for Gr-4D. A) RT-qPCR analysis showing knockdown of *NRF2* expression by more than 95% in WI-38 cells transfected with siRNA oligos (normalized to *PPIA*, n = 8 each; one-way ANOVA, followed by Dunnett’s multiple comparison, *P* < 0.0001). B) WI-38 cells were transfected with control or *NRF2* siRNA and then treated with Gr-4D (from two different vendors) at 3 different doses (2.5/3.75/5 μM), before subject to H_2_O_2_ or no H_2_O_2_ treatment, and scored for propidium iodide staining. Note that Gr-4D (from two different vendors) significantly increased the percentage of PI-positive cells, particularly, in *NRF2*-deficient WI-38 cells at higher doses in the absence of H_2_O_2_. Hence, we treated cells with a lower dose of H_2_O_2_ (500uM instead of 700uM) to prevent ceiling of death for *NRF2*-deficent cells. Gr-4D also protected *NRF2(-)* cells from H_2_O_2_, just like what it did in *NRF2(+)* cells. B & C) In parallel to our RNA-seq or microarray analysis of WI-38 treated with small molecules (in the absence of H_2_O_2_), cells were incubated with these molecules (Gr-4D, 2.5/3.75/5 μM, n = 8 for RNA-seq; or others, 10 μM, n = 3 for microarrays) or DMSO control (0.1%, n = 8 for RNA-seq or 6-8 for microarray) for 24 hours in 96-well plates and then analyzed for cell viability upon 3 hours of H_2_O_2_ treatment. Shown is the percentage of cells that scored positively for propidium iodide staining following treatment with DMSO control or small molecules (named according to the source of vendors: CB, ChemBridge; CD, ChemDiv; MP, MolPort; and V, Vitascreen). Student’s t-test, * *P* < 0.05; ** *P* < 0.01; *** *P* < 0.001; n.s. not significant; normalized to respective controls (color-coded).

Supplemental Figure 7 (related to Figure 3). Pearson’s correlation between global transcriptional profiles for cells treated with small molecules. Shown is the normalized expression of the 18,683 probe sets detected in every array, grouped by unsupervised clustering using Pearson correlation coefficient as a distance metric. Note that most DMSO (0.1%) controls were clustered together on the tree, as were the six rapamycin (2.5 μM)-treated samples (highlighted in blue). The molecules were named according to the source of vendors (CB, ChemBridge; CD, ChemDiv; V, Vitascreen), and Gr-4A molecules were obtained from ChemDiv and Asinex.

Supplemental Figure 8 (related to Figure 2). Inhibitory effects of certain small molecules on PARP. Two known PARP inhibitors, PJ-34 (Tocris) (IC_50_: ∼20 nM) and MK-4827 (IC_50_: ∼3.8 nM), were shown to inhibit human PARP1 (at 200 nM and 40 nM, respectively), as indicated by a substantial increase of normalized fluorescence (n = 2). 10 of the 51 repurchased molecules also inhibited PARP at 10 μM. Note that all the molecules from group 2 assayed (one was not available for repurchasing), plus two group-8 molecules (analogs of the 4-amino-1,8-naphthalimide PARP inhibitor), were confirmed to be PARP inhibitors in this assay.

Supplemental Figure 9 (related to Table 1). Protective effects of certain small molecules against poly-Q toxicity. 51 repurchased molecules were introduced initially at 10 μM to neuron-like PC12 cells that express poly(Q)-tagged GFP (Q103-Htt-EGFP), and candidates showing protective effects were further retested at multiple doses (2.5 μM, 5 μM, 10 μM and 20 μM) to analyze their effects on ATP content upon the induction of toxic poly(Q)103-Htt-EGFP aggregates. The parental PC12 cells (WT) that do not express poly(Q) were used as the control to demonstrate the specificity of protective effects. Note that 48 hours induction of poly(Q)103-Htt-EGFP reduced ATP content substantially (right, bottom panel), and several small molecules produced modest yet significant effects to enhance ATP content. However, except for Gr-6B, under non-induced conditions, these small molecules actually exerted cell toxicity and reduced ATP content in 72 hours (n = 6. Student’s t-test, * *P* < 0.05; ** *P* < 0.01; *** *P* < 0.001).

Supplemental Figure 10 (related to Figure 4). Effects of Gr-4D on the expression of reporters for several pathways known to influence longevity in *C. elegans*. L4 animals expressing different *gfp* fusions were raised on plates containing DMSO (as the control) or different compounds as indicated for ∼30 hours and then imaged for analysis. At least 12-15 worms were analyzed and shown are representative images. Note that Gr-4D (from two different suppliers), at a dose that extended lifespan, caused modest induction of the *gst-4* reporter, but not the others. No obvious effects of Gr-4D were observed on the number of LGG-1::GFP puncta, which was usually used as the readout of perturbed autophagy (data not shown).

## Supplemental Table Legends

Supplemental Table 1 (related to Table 1). Summary of 61 primary hits that promote resistance to H_2_O_2_. 1^st^ tab: 209 primary screen hits; 2^nd^ tab: 61 primary hits; 3^rd^ tab: 32 “core-set” hits. Potential “pan-assay interference compounds” (PAINS) are indicated. Dose-response curves and derived EC_50_ values are shown for the library compounds identified. Note that significant reduction of performance (fold change < 1.50, highlighted in red) relative to the initial score, likely due to compromised stability, was observed in certain cases. This table also shows data for repurchased small molecules in the follow-up characterizations, including validation retests and other cell-based assays, as well as multiple independent lifespan assays performed in *C. elegans*. For lifespan assays using wild type, FuDR (100 μM) was applied to block progeny production, or temperature-sensitive *fer-15; fem-1* sterile mutants were used instead (see Supplemental Table 8 for details).

Supplemental Table 2 (related to Table 1). 1^st^ tab: Cell death-imaging analysis of 32 core-set small molecules. Consistent with ATP assay results to assess cell viability, the 32 core-set small molecules also reduced the fraction of propidium iodide-positive dying/dead WI-38 cells upon H_2_O_2_ (n = 6 for each molecule, Student’s t-test, three consecutive time points – 3, 4 and 5 hours of H_2_O_2_ incubation) [Compare the percentage of dead cells in the presence of the small molecules (green, decreased; red, increased) with the reference percentage in the DMSO control (shown at the top)]. Shown are representative data for at least three independent experiments. Note that besides the 32 core-set molecules, three PARP inhibitors (*, as reference) only rendered WI-38 cells very modest protection. 2^nd^ tab: Likewise, in parallel to assays of WI-38 cells, 22 small molecules also protected primary, non-transformed human dermal fibroblasts (HDFs, from multiple donors) from H_2_O_2_ (2 trials). Fold change less than 1.5 is highlighted in red.

Supplemental Table 3 (related to Table 1). Protection of WI-38 cells from the heavy metal CdCl_2_ by small molecules. Among molecules retested, at least 17 increased the resistance of WI-38 cells to both H_2_O_2_ and cadmium (in two independent experiments). Fold changes less than 1.5 are highlighted in red. In a few cases (Gr-7A, Gr-7B, Gr-9A and Gr-9B), we did not observe H_2_O_2_ resistance, potentially due to chemical instability (not shown). Thus, we were unable to judge these molecules for cadmium resistance.

Supplemental Table 4 (related to Table 1). Long-term effects of small molecules on ATP levels and DNA damage-associated markers in the absence of H_2_O_2_. 1^st^ tab: included are the data shown in Supplemental Figure 5, which addressed long-term effects of small molecules on ATP levels. Of 32 core-set hits, 28 molecules were analyzed in multiple 384-well plates in replicates in two batches. 11 small molecules that decreased the ATP content by 50% or more on day 5 of treatment, due to their effects reducing cell growth and/or cell viability, are highlighted. Unlike rapamycin, which also reduced ATP levels substantially, 9 of these 11 molecules (Gr-3B, Gr-6C, Gr-7A, Gr-7B, Gr-7C, Gr-9B, O11, O21 & O27) also produced cell toxicity (by the examination of cell morphology, data not shown). The other two, Gr-1F and Gr-3C, caused DNA damages, like the others. 2^nd^ tab: WI-38 cells that had been treated with small molecules (10 μM, 24 hours) were analyzed in two independent experiments by immuno-staining for two DNA damage-associated markers, phosphorylated-*γ*H2A.X and phosphorylated-TP53BP1. Small molecules that induced both markers (in the absence of H_2_O_2_) were scored positively for DNA-damaging. Normalized values for marker-positive cell fractions are shown (n = 3, Student’ t-test).

In experiment 2 to analyze DNA damage-associated markers, H_2_O_2_ was also introduced during the last 3 hours of incubation for one set of plates. For the other set that was not exposed to H_2_O_2_, several wells of cells were treated with the controls, including DMSO (0.1%) and doxorubicin (300 nM) for 24 hours or for H_2_O_2_ (700 μM) 3 hours. The fractions of DNA damage marker-positive cells were: for *γ*H2A.X-P – DMSO negative control, ∼1.7±0.4% (n = 6); doxorubicin positive control, ∼16.9±2.1% (n = 6); and H_2_O_2_ positive control, ∼50.9±4.2% (n = 3). For TP53BP1-P – DMSO negative control, ∼1.0±0.2% (n = 6); doxorubicin positive control, ∼15.1±1.4% (n = 6); and H_2_O_2_ positive control, ∼45.8±5.1% (n = 3). Green, decreased; red, increased. In certain cases, we observed a strong trend of increasing DNA damage-associated markers, while the p values did not reach statistical significance, possibly due to a rather large variation of the samples. Note that besides the 32 core-set molecules, as expected, three PARP inhibitors (*) increased significantly the fraction of cells positive for DNA damage-associated markers upon H_2_O_2_ treatment.

Supplemental Table 5 (related to Figure 3). A) List of significant genes identified in RNA-seq analysis for cells treated with Gr-4D. WI-38 cells were treated with Gr-4D (2.5 μM, n = 6; from two different vendors – CB, ChemBridge; MP, MolPort) for 24 hours (in the absence of H_2_O_2_) and then analyzed by RNA-seq to address Gr-4D’s effects on gene expression. Genes whose expression levels are either up- or down-regulated significantly (FDR-adjusted q value < 0.05) are shown in red or blue, respectively (unlogged effect size > 1.5-fold, or > 1.25-fold, plus significant genes identified in samples treated with both Gr-4Ds). Total number of significant genes is indicated on the top. B) List of significant genes identified by SAM analysis of microarray for cells treated with other small molecules. WI-38 cells were treated with small molecules (10 μM, unless otherwise specified, n = 3) for 24 hours (in the absence of H_2_O_2_) and then analyzed by microarray analysis to address small molecules’ effects on gene expression. Besides the 0.1% DMSO negative controls (n = 15), rapamycin (2.5 μM, n = 6) was included for comparison as well. For each small molecule treatment (group-4 and group-5 molecules from different vendors: CB, ChemBridge; CD, ChemDiv; and V, Vitascreen), genes whose expression levels are either up- or down-regulated significantly are shown in red or blue, respectively. Total number of significant genes is indicated on the top, with unlogged fold change (>= 1.5) and FDR-adjusted q value [< 0.10 (10%)] shown. Overlap between significant genes affected by the same molecules (from different vendors) or molecules of the same structural group were shown as well.

Supplemental Table 6 (related to Figure 3). A) Summary of pathway analysis of Gr-4D RNA-seq data. 1^st^ tab: analysis of 550 significant genes (normalized fold change > = 1.25) using Enrichr or PANTHER GO analysis tool. Shown were top pathways: for Enrichr-KEGG analysis, with a high combined score (>= 10), which combines both FDR-adjusted P value and Z score and represents the best rankings of enrichment (Chen et al., 2013); for PANTHER GO analysis, ranked by FDR-adjusted significance (< 0.05) for each GO term category (pathways, molecular function, biological process, etc). Significant genes (up- or down-regulated, in red or blue) were shown also for several top pathways of interest. 2^nd^ tab: comparison of significant genes identified in WI-38 cells treated with our small molecules with other perturbation conditions (gene knockdown or small-molecule treatment). B) Summary of pathway analysis results for other small molecules analyzed by microarray. One tab showed only top pathways (with FDR-adjusted significance < 0.05), plus genes of the pathway indicated by certain analysis for each molecule, and another tab showed the iLINCS similarity analysis results. Note that no significant similarity was suggested using significant genes for Gr-5D.

Supplemental Table 7 (related to Table 1). Potential mTOR inhibition by certain small molecules. Small molecules were examined initially at 4 doses (2.5 μM, 5 μM, 10 μM and 20 μM, n = 4 for each dose) and then retested twice by In-Cell Western analyses for their effects on the phosphorylation status of ribosomal protein S6, a readout of mTOR activity. Small molecules that caused more than 30% reduction in the p-RPS6/RPS6 ratio in at least 2 of 3 independent experiments were regarded as candidate mTOR inhibitors. Note that rapamycin, as the positive control, reduced the normalized ratio of p-RPS6/RPS6 substantially. Unlike rapamycin, these putative upstream inhibitors were toxic to cells at the doses examined.

Supplemental Table 8 (related to Figure 4). Effects of small molecules on *C. elegans’* lifespan and H_2_O_2_-resistance. Multiple independent trials were conducted to analyze the effects of small molecules on lifespan. Trials 1 and 2 were conducted in liquid (1.0X10^9 bacteria/ml and 2.5X10^9 bacteria/ml, respectively), using wild-type animals in the presence of 100 μM FuDR to block progeny production; and small molecules were administered at the final concentration of 60 μM (containing 0.3% DMSO in liquid). Trials 3 and 4 were conducted on plates (2.0X10^10 bacteria/ml and 1.0X10^11 bacteria/ml, respectively; 100 μl per plate, treated with UV and kanamycin), using temperature-sensitive sterile mutant animals in the absence of FuDR; and small molecules were provided at 60 μM (containing 0.2% DMSO on plate). Two confirmative trials 5 and 6 were also conducted, using the same conditions as trials 3 and 4 (only results for 4 molecules are shown). For the H_2_O_2_-stress resistance assays, wild-type animals were treated first with small molecules (60 μM) and then subjected to 0.5 mM H_2_O_2_ in the presence of small molecules (1^st^ trial, 8-day-old; 2^nd^ trial, 6-day-old. Note that the mean survival times of controls are different between these two trials). Shown are the normalized changes of mean survival time (mean ± SEM; in days for lifespan assay, and in hours for H_2_O_2_ stress assay) and observed/total numbers of animals (log-rank test, *P* < 0.05 high-lighted). Green, increased; red, decreased.

Supplemental Table 9. Potential targets for small molecules identified in other assays. Shown are potential (direct or indirect) human protein targets for 9 small molecules that were identified and reported previously in other screens. 9 of our small-molecule hits scored positively (from PubChem, https://www.ncbi.nlm.nih.gov/pccompound). 1^st^ tab: list of potential (direct or indirect) human protein targets of each small molecule (see URLs below), based on these screens.

## References

Aiken, C.T., Tobin, A.J., and Schweitzer, E.S. (2004). A cell-based screen for drugs to treat Huntington’s disease. Neurobiol Dis 16, 546–555.

Alavez, S., Vantipalli, M.C., Zucker, D.J., Klang, I.M., and Lithgow, G.J. (2011). Amyloid-binding compounds maintain protein homeostasis during ageing and extend lifespan. Nature 472, 226–229.

Anand, P., Sundaram, C., Jhurani, S., Kunnumakkara, A.B., and Aggarwal, B.B. (2008). Curcumin and cancer: an “old-age” disease with an “age-old” solution. Cancer Lett 267, 133–164.

Aso, E., and Ferrer, I. (2014). Cannabinoids for treatment of Alzheimer’s disease: moving toward the clinic. Front Pharmacol 5, 37.

Bae, S.H., Sung, S.H., Oh, S.Y., Lim, J.M., Lee, S.K., Park, Y.N., Lee, H.E., Kang, D., and Rhee, S.G. (2013). Sestrins activate Nrf2 by promoting p62-dependent autophagic degradation of Keap1 and prevent oxidative liver damage. Cell Metab 17, 73–84.

Baell, J., and Walters, M.A. (2014). Chemistry: Chemical con artists foil drug discovery. Nature 513, 481–483.

Baell, J.B., and Holloway, G.A. (2010). New substructure filters for removal of pan assay interference compounds (PAINS) from screening libraries and for their exclusion in bioassays. J Med Chem 53, 2719–2740.

Bai, P., Canto, C., Oudart, H., Brunyanszki, A., Cen, Y., Thomas, C., Yamamoto, H., Huber, A., Kiss, B., Houtkooper, R.H., et al. (2011). PARP-1 inhibition increases mitochondrial metabolism through SIRT1 activation. Cell Metab 13, 461–468.

Banasik, M., Komura, H., Shimoyama, M., and Ueda, K. (1992). Specific inhibitors of poly(ADP-ribose) synthetase and mono(ADP-ribosyl)transferase. J Biol Chem 267, 1569–1575.

Bartke, A. (2011). Single-gene mutations and healthy ageing in mammals. Philos Trans R Soc Lond B Biol Sci 366, 28–34.

Batovska, D.I., and Todorova, I.T. (2010). Trends in utilization of the pharmacological potential of chalcones. Curr Clin Pharmacol 5, 1–29.

Benjamin, D., Colombi, M., Moroni, C., and Hall, M.N. (2011). Rapamycin passes the torch: a new generation of mTOR inhibitors. Nat Rev Drug Discov 10, 868–880.

Birse, R.T., Choi, J., Reardon, K., Rodriguez, J., Graham, S., Diop, S., Ocorr, K., Bodmer, R., and Oldham, S. (2010). High-fat-diet-induced obesity and heart dysfunction are regulated by the TOR pathway in Drosophila. Cell Metab 12, 533–544.

Bishop, N.A., and Guarente, L. (2007). Two neurons mediate diet-restriction-induced longevity in C. elegans. Nature 447, 545–549.

Bjedov, I., Toivonen, J.M., Kerr, F., Slack, C., Jacobson, J., Foley, A., and Partridge, L. (2010). Mechanisms of life span extension by rapamycin in the fruit fly Drosophila melanogaster. Cell Metab 11, 35–46.

Blackwell, T.K., Steinbaugh, M.J., Hourihan, J.M., Ewald, C.Y., and Isik, M. (2015). SKN-1/Nrf, stress responses, and aging in Caenorhabditis elegans. Free Radic Biol Med 88, 290–301.

Booz, G.W. (2011). Cannabidiol as an emergent therapeutic strategy for lessening the impact of inflammation on oxidative stress. Free Radic Biol Med 51, 1054–1061.

Borst, P., Evers, R., Kool, M., and Wijnholds, J. (1999). The multidrug resistance protein family. Biochim Biophys Acta 1461, 347–357.

Brasnyo, P., Molnar, G.A., Mohas, M., Marko, L., Laczy, B., Cseh, J., Mikolas, E., Szijarto, I.A., Merei, A., Halmai, R., et al. (2011). Resveratrol improves insulin sensitivity, reduces oxidative stress and activates the Akt pathway in type 2 diabetic patients. Br J Nutr 106, 383–389.

Budanov, A.V., and Karin, M. (2008). p53 target genes sestrin1 and sestrin2 connect genotoxic stress and mTOR signaling. Cell 134, 451–460.

Budanov, A.V., Sablina, A.A., Feinstein, E., Koonin, E.V., and Chumakov, P.M. (2004). Regeneration of peroxiredoxins by p53-regulated sestrins, homologs of bacterial AhpD. Science 304, 596–600.

Cabreiro, F., Au, C., Leung, K.Y., Vergara-Irigaray, N., Cocheme, H.M., Noori, T., Weinkove, D., Schuster, E., Greene, N.D., and Gems, D. (2013). Metformin retards aging in C. elegans by altering microbial folate and methionine metabolism. Cell 153, 228–239.

Calamini, B., Silva, M.C., Madoux, F., Hutt, D.M., Khanna, S., Chalfant, M.A., Saldanha, S.A., Hodder, P., Tait, B.D., Garza, D., et al. (2011). Small-molecule proteostasis regulators for protein conformational diseases. Nat Chem Biol 8, 185–196.

Cao, S.Q., Xu, Q.T., Zhou, H.J., Cao, Y.J., Zhu, Y., Yu, F., and Kuai, B.K. (2003). [Screening for lifespan-extension mutants with paraquat in Arabidopsis]. Shi Yan Sheng Wu Xue Bao 36, 233–237.

Carmona-Gutierrez, D., Zimmermann, A., Kainz, K., Pietrocola, F., Chen, G., Maglioni, S., Schiavi, A., Nah, J., Mertel, S., Beuschel, C.B., et al. (2019). The flavonoid 4,4’-dimethoxychalcone promotes autophagy-dependent longevity across species. Nat Commun 10, 651.

Caughey, B., and Lansbury, P.T. (2003). Protofibrils, pores, fibrils, and neurodegeneration: separating the responsible protein aggregates from the innocent bystanders. Annu Rev Neurosci 26, 267–298.

Chan, N., Pires, I.M., Bencokova, Z., Coackley, C., Luoto, K.R., Bhogal, N., Lakshman, M., Gottipati, P., Oliver, F.J., Helleday, T., et al. (2010). Contextual synthetic lethality of cancer cell kill based on the tumor microenvironment. Cancer Res 70, 8045–8054.

Chen, E.Y., Tan, C.M., Kou, Y., Duan, Q., Wang, Z., Meirelles, G.V., Clark, N.R., and Ma’ayan, A. (2013). Enrichr: interactive and collaborative HTML5 gene list enrichment analysis tool. BMC Bioinformatics 14, 128.

Cheng, C.W., Adams, G.B., Perin, L., Wei, M., Zhou, X., Lam, B.S., Da Sacco, S., Mirisola, M., Quinn, D.I., Dorff, T.B., et al. (2014). Prolonged fasting reduces IGF-1/PKA to promote hematopoietic-stem-cell-based regeneration and reverse immunosuppression. Cell Stem Cell 14, 810–823.

Cohen, E., Paulsson, J.F., Blinder, P., Burstyn-Cohen, T., Du, D., Estepa, G., Adame, A., Pham, H.M., Holzenberger, M., Kelly, J.W., et al. (2009). Reduced IGF-1 signaling delays age-associated proteotoxicity in mice. Cell 139, 1157–1169.

Costantino, G., Macchiarulo, A., Camaioni, E., and Pellicciari, R. (2001). Modeling of poly(ADP-ribose)polymerase (PARP) inhibitors. Docking of ligands and quantitative structure-activity relationship analysis. J Med Chem 44, 3786–3794.

Dazert, E., and Hall, M.N. (2011). mTOR signaling in disease. Curr Opin Cell Biol 23, 744–755.

de Castro, E., Hegi de Castro, S., and Johnson, T.E. (2004). Isolation of long-lived mutants in Caenorhabditis elegans using selection for resistance to juglone. Free Radic Biol Med 37, 139–145.

Ding, A.J., Zheng, S.Q., Huang, X.B., Xing, T.K., Wu, G.S., Sun, H.Y., Qi, S.H., and Luo, H.R. (2017). Current Perspective in the Discovery of Anti-aging Agents from Natural Products. Nat Prod Bioprospect 7, 335–404.

Ekins, S., Nikolsky, Y., Bugrim, A., Kirillov, E., and Nikolskaya, T. (2007). Pathway mapping tools for analysis of high content data. Methods Mol Biol 356, 319–350.

Fong, P.C., Boss, D.S., Yap, T.A., Tutt, A., Wu, P., Mergui-Roelvink, M., Mortimer, P., Swaisland, H., Lau, A., O’Connor, M.J., et al. (2009). Inhibition of poly(ADP-ribose) polymerase in tumors from BRCA mutation carriers. N Engl J Med 361, 123–134.

Fontana, L., Partridge, L., and Longo, V.D. (2010). Extending healthy life span--from yeast to humans. Science 328, 321–326.

Geiss-Friedlander, R., and Melchior, F. (2007). Concepts in sumoylation: a decade on. Nat Rev Mol Cell Biol 8, 947–956.

Giannakou, M.E., Goss, M., Junger, M.A., Hafen, E., Leevers, S.J., and Partridge, L. (2004). Long-lived Drosophila with overexpressed dFOXO in adult fat body. Science 305, 361.

Guevara-Aguirre, J., Balasubramanian, P., Guevara-Aguirre, M., Wei, M., Madia, F., Cheng, C.W., Hwang, D., Martin-Montalvo, A., Saavedra, J., Ingles, S., et al. (2011). Growth hormone receptor deficiency is associated with a major reduction in pro-aging signaling, cancer, and diabetes in humans. Sci Transl Med 3, 70ra13.

Harper, J.M., Salmon, A.B., Leiser, S.F., Galecki, A.T., and Miller, R.A. (2007). Skin-derived fibroblasts from long-lived species are resistant to some, but not all, lethal stresses and to the mitochondrial inhibitor rotenone. Aging Cell 6, 1–13.

Harper, J.M., Wang, M., Galecki, A.T., Ro, J., Williams, J.B., and Miller, R.A. (2011). Fibroblasts from long-lived bird species are resistant to multiple forms of stress. The Journal of experimental biology 214, 1902–1910.

Harrington, S.C., Simari, R.D., and Conover, C.A. (2007). Genetic deletion of pregnancy-associated plasma protein-A is associated with resistance to atherosclerotic lesion development in apolipoprotein E-deficient mice challenged with a high-fat diet. Circ Res 100, 1696–1702.

Harrison, D.E., Strong, R., Sharp, Z.D., Nelson, J.F., Astle, C.M., Flurkey, K., Nadon, N.L., Wilkinson, J.E., Frenkel, K., Carter, C.S., et al. (2009). Rapamycin fed late in life extends lifespan in genetically heterogeneous mice. Nature 460, 392–395.

Hayden, M.S., and Ghosh, S. (2012). NF-kappaB, the first quarter-century: remarkable progress and outstanding questions. Genes Dev 26, 203–234.

Hickey, C.M., Wilson, N.R., and Hochstrasser, M. (2012). Function and regulation of SUMO proteases. Nat Rev Mol Cell Biol 13, 755–766.

Hubbard, B.P., and Sinclair, D.A. (2014). Small molecule SIRT1 activators for the treatment of aging and age-related diseases. Trends Pharmacol Sci 35, 146–154.

Hur, W., and Gray, N.S. (2011). Small molecule modulators of antioxidant response pathway. Curr Opin Chem Biol 15, 162–173.

Hwangbo, D.S., Gershman, B., Tu, M.P., Palmer, M., and Tatar, M. (2004). Drosophila dFOXO controls lifespan and regulates insulin signalling in brain and fat body. Nature 429, 562–566.

Hybertson, B.M., Gao, B., Bose, S.K., and McCord, J.M. (2011). Oxidative stress in health and disease: the therapeutic potential of Nrf2 activation. Mol Aspects Med 32, 234–246.

Ikeno, Y., Hubbard, G.B., Lee, S., Cortez, L.A., Lew, C.M., Webb, C.R., Berryman, D.E., List, E.O., Kopchick, J.J., and Bartke, A. (2009). Reduced incidence and delayed occurrence of fatal neoplastic diseases in growth hormone receptor/binding protein knockout mice. J Gerontol A Biol Sci Med Sci 64, 522–529.

Inoki, K., Zhu, T., and Guan, K.L. (2003). TSC2 mediates cellular energy response to control cell growth and survival. Cell 115, 577–590.

Irwin, J.J., Sterling, T., Mysinger, M.M., Bolstad, E.S., and Coleman, R.G. (2012). ZINC: a free tool to discover chemistry for biology. J Chem Inf Model 52, 1757–1768.

Jenny, N.S. (2012). Inflammation in Aging: Cause, Effect, or Both? Discov Med 73, 451–460.

Joshi, P.K., Pirastu, N., Kentistou, K.A., Fischer, K., Hofer, E., Schraut, K.E., Clark, D.W., Nutile, T., Barnes, C.L.K., Timmers, P., et al. (2017). Genome-wide meta-analysis associates HLA-DQA1/DRB1 and LPA and lifestyle factors with human longevity. Nat Commun 8, 910.

Kalaany, N.Y., and Sabatini, D.M. (2009). Tumours with PI3K activation are resistant to dietary restriction. Nature 458, 725–731.

Kalender, A., Selvaraj, A., Kim, S.Y., Gulati, P., Brule, S., Viollet, B., Kemp, B.E., Bardeesy, N., Dennis, P., Schlager, J.J., et al. (2010). Metformin, independent of AMPK, inhibits mTORC1 in a rag GTPase-dependent manner. Cell Metab 11, 390–401.

Kaletsky, R., Lakhina, V., Arey, R., Williams, A., Landis, J., Ashraf, J., and Murphy, C.T. (2016). The C. elegans adult neuronal IIS/FOXO transcriptome reveals adult phenotype regulators. Nature 529, 92–96.

Kapeta, S., Chondrogianni, N., and Gonos, E.S. (2010). Nuclear erythroid factor 2-mediated proteasome activation delays senescence in human fibroblasts. J Biol Chem 285, 8171–8184.

Keiser, M.J., Roth, B.L., Armbruster, B.N., Ernsberger, P., Irwin, J.J., and Shoichet, B.K. (2007). Relating protein pharmacology by ligand chemistry. Nat Biotechnol 25, 197–206.

Kennedy, B.K., Austriaco, N.R., Jr., Zhang, J., and Guarente, L. (1995). Mutation in the silencing gene SIR4 can delay aging in S. cerevisiae. Cell 80, 485–496.

Kennedy, M.A., Rakoczy, S.G., and Brown-Borg, H.M. (2003). Long-living Ames dwarf mouse hepatocytes readily undergo apoptosis. Exp Gerontol 38, 997–1008.

Kenyon, C., Chang, J., Gensch, E., Rudner, A., and Tabtiang, R. (1993). A C. elegans mutant that lives twice as long as wild type. Nature 366, 461–464.

Kenyon, C.J. (2010a). The genetics of ageing. In Nature, pp. 504–512.

Kenyon, C.J. (2010b). The genetics of ageing. Nature 464, 504–512.

Kim, Y., and Sun, H. (2007). Functional genomic approach to identify novel genes involved in the regulation of oxidative stress resistance and animal lifespan. Aging Cell 6, 489–503.

Kolovou, V., Bilianou, H., Giannakopoulou, V., Kalogeropoulos, P., Mihas, C., Kouris, M., Cokkinos, D.V., Boutsikou, M., Hoursalas, I., Mavrogeni, S., et al. (2017). Five gene variants in nonagenarians, centenarians and average individuals. Arch Med Sci 13, 1130–1141.

Kovacs, J.J., Murphy, P.J., Gaillard, S., Zhao, X., Wu, J.T., Nicchitta, C.V., Yoshida, M., Toft, D.O., Pratt, W.B., and Yao, T.P. (2005). HDAC6 regulates Hsp90 acetylation and chaperone-dependent activation of glucocorticoid receptor. Mol Cell 18, 601–607.

Kumar, A., and Zhang, K.Y. (2015). Advances in the development of SUMO specific protease (SENP) inhibitors. Comput Struct Biotechnol J 13, 204–211.

Langley, B., Gensert, J.M., Beal, M.F., and Ratan, R.R. (2005). Remodeling chromatin and stress resistance in the central nervous system: histone deacetylase inhibitors as novel and broadly effective neuroprotective agents. Curr Drug Targets CNS Neurol Disord 4, 41–50.

Lawrence, T. (2009). The nuclear factor NF-kappaB pathway in inflammation. Cold Spring Harb Perspect Biol 1, a001651.

Le Couteur, D.G., McLachlan, A.J., Quinn, R.J., Simpson, S.J., and de Cabo, R. (2012). Aging biology and novel targets for drug discovery. J Gerontol A Biol Sci Med Sci 67, 168–174.

Lee, J.H., Budanov, A.V., and Karin, M. (2013). Sestrins orchestrate cellular metabolism to attenuate aging. Cell Metab 18, 792–801.

Lee, J.H., Budanov, A.V., Park, E.J., Birse, R., Kim, T.E., Perkins, G.A., Ocorr, K., Ellisman, M.H., Bodmer, R., Bier, E., et al. (2010). Sestrin as a feedback inhibitor of TOR that prevents age-related pathologies. Science 327, 1223–1228.

Lee, J.H., Budanov, A.V., Talukdar, S., Park, E.J., Park, H.L., Park, H.W., Bandyopadhyay, G., Li, N., Aghajan, M., Jang, I., et al. (2012). Maintenance of metabolic homeostasis by Sestrin2 and Sestrin3. Cell Metab 16, 311–321.

Leiser, S.F., and Miller, R.A. (2010). Nrf2 signaling, a mechanism for cellular stress resistance in long-lived mice. Mol Cell Biol 30, 871–884.

Lerner, C., Bitto, A., Pulliam, D., Nacarelli, T., Konigsberg, M., Van Remmen, H., Torres, C., and Sell, C. (2013). Reduced mammalian target of rapamycin activity facilitates mitochondrial retrograde signaling and increases life span in normal human fibroblasts. Aging Cell 12, 966–977.

Lewis, K.N., Mele, J., Hornsby, P.J., and Buffenstein, R. (2012). Stress resistance in the naked mole-rat: the bare essentials - a mini-review. Gerontology 58, 453–462.

Lewis, K.N., Wason, E., Edrey, Y.H., Kristan, D.M., Nevo, E., and Buffenstein, R. (2015). Regulation of Nrf2 signaling and longevity in naturally long-lived rodents. Proc Natl Acad Sci U S A 112, 3722–3727.

Li, W., Li, X., and Miller, R.A. (2014). ATF4 activity: a common feature shared by many kinds of slow-aging mice. Aging Cell 13, 1012–1018.

Libina, N., Berman, J.R., and Kenyon, C. (2003). Tissue-specific activities of C. elegans DAF-16 in the regulation of lifespan. Cell 115, 489–502.

Luo, X., and Kraus, W.L. (2012). On PAR with PARP: cellular stress signaling through poly(ADP-ribose) and PARP-1. Genes Dev 26, 417–432.

Magesh, S., Chen, Y., and Hu, L. (2012). Small molecule modulators of Keap1-Nrf2-ARE pathway as potential preventive and therapeutic agents. Med Res Rev 32, 687–726.

Marchand, A., Tomkiewicz, C., Magne, L., Barouki, R., and Garlatti, M. (2006). Endoplasmic reticulum stress induction of insulin-like growth factor-binding protein-1 involves ATF4. J Biol Chem 281, 19124–19133.

Martin-Montalvo, A., Mercken, E.M., Mitchell, S.J., Palacios, H.H., Mote, P.L., Scheibye-Knudsen, M., Gomes, A.P., Ward, T.M., Minor, R.K., Blouin, M.J., et al. (2013). Metformin improves healthspan and lifespan in mice. Nat Commun 4, 2192.

Mason, K.A., Raju, U., Buchholz, T.A., Wang, L., Milas, Z.L., and Milas, L. (2014). Poly (ADP-ribose) polymerase inhibitors in cancer treatment. Am J Clin Oncol 37, 90–100.

Miller, R.A. (2009). Cell stress and aging: new emphasis on multiplex resistance mechanisms. J Gerontol A Biol Sci Med Sci 64, 179–182.

Morley, J.F., Brignull, H.R., Weyers, J.J., and Morimoto, R.I. (2002). The threshold for polyglutamine-expansion protein aggregation and cellular toxicity is dynamic and influenced by aging in Caenorhabditis elegans. Proc Natl Acad Sci U S A 99, 10417–10422.

Mouchiroud, L., Houtkooper, R.H., Moullan, N., Katsyuba, E., Ryu, D., Canto, C., Mottis, A., Jo, Y.S., Viswanathan, M., Schoonjans, K., et al. (2013). The NAD(+)/Sirtuin Pathway Modulates Longevity through Activation of Mitochondrial UPR and FOXO Signaling. Cell 154, 430–441.

Murphy, C.T., McCarroll, S.A., Bargmann, C.I., Fraser, A., Kamath, R.S., Ahringer, J., Li, H., and Kenyon, C. (2003). Genes that act downstream of DAF-16 to influence the lifespan of Caenorhabditis elegans. Nature 424, 277–283.

Nogueira, V., Park, Y., Chen, C.C., Xu, P.Z., Chen, M.L., Tonic, I., Unterman, T., and Hay, N. (2008). Akt determines replicative senescence and oxidative or oncogenic premature senescence and sensitizes cells to oxidative apoptosis. Cancer Cell 14, 458–470.

Ogg, S., Paradis, S., Gottlieb, S., Patterson, G.I., Lee, L., Tissenbaum, H.A., and Ruvkun, G. (1997). The Fork head transcription factor DAF-16 transduces insulin-like metabolic and longevity signals in C. elegans. Nature 389, 994–999.

Panier, S., and Boulton, S.J. (2014). Double-strand break repair: 53BP1 comes into focus. Nat Rev Mol Cell Biol 15, 7–18.

Perez, V.I., Bokov, A., Van Remmen, H., Mele, J., Ran, Q., Ikeno, Y., and Richardson, A. (2009). Is the oxidative stress theory of aging dead? Biochim Biophys Acta 1790, 1005–1014.

Perez-Cadahia, B., Drobic, B., Khan, P., Shivashankar, C.C., and Davie, J.R. (2010). Current understanding and importance of histone phosphorylation in regulating chromatin biology. Curr Opin Drug Discov Devel 13, 613–622.

Pickering, A.M., Linder, R.A., Zhang, H., Forman, H.J., and Davies, K.J. (2012). Nrf2-dependent induction of proteasome and Pa28alphabeta regulator are required for adaptation to oxidative stress. J Biol Chem 287, 10021–10031.

Pinkston, J.M., Garigan, D., Hansen, M., and Kenyon, C. (2006). Mutations that increase the life span of C. elegans inhibit tumor growth. Science 313, 971–975.

Pirinen, E., Canto, C., Jo, Y.S., Morato, L., Zhang, H., Menzies, K.J., Williams, E.G., Mouchiroud, L., Moullan, N., Hagberg, C., et al. (2014). Pharmacological Inhibition of poly(ADP-ribose) polymerases improves fitness and mitochondrial function in skeletal muscle. Cell Metab 19, 1034–1041.

Prodromou, C. (2016). Mechanisms of Hsp90 regulation. Biochem J 473, 2439–2452.

Robida-Stubbs, S., Glover-Cutter, K., Lamming, D.W., Mizunuma, M., Narasimhan, S.D., Neumann-Haefelin, E., Sabatini, D.M., and Blackwell, T.K. (2012). TOR signaling and rapamycin influence longevity by regulating SKN-1/Nrf and DAF-16/FoxO. Cell Metab 15, 713–724.

Rohn, T.T. (2010). The role of caspases in Alzheimer’s disease; potential novel therapeutic opportunities. Apoptosis 15, 1403–1409.

Ross, C.A., and Poirier, M.A. (2004). Protein aggregation and neurodegenerative disease. Nat Med 10 Suppl, S10–17.

Rouleau, M., Patel, A., Hendzel, M.J., Kaufmann, S.H., and Poirier, G.G. (2010). PARP inhibition: PARP1 and beyond. Nat Rev Cancer 10, 293–301.

Sahni, S.K., Rydkina, E., and Sahni, A. (2008). The proteasome inhibitor MG132 induces nuclear translocation of erythroid transcription factor Nrf2 and cyclooxygenase-2 expression in human vascular endothelial cells. Thromb Res 122, 820–825.

Sahu, N.K., Balbhadra, S.S., Choudhary, J., and Kohli, D.V. (2012). Exploring pharmacological significance of chalcone scaffold: a review. Curr Med Chem 19, 209–225.

Salminen, A., Huuskonen, J., Ojala, J., Kauppinen, A., Kaarniranta, K., and Suuronen, T. (2008). Activation of innate immunity system during aging: NF-kB signaling is the molecular culprit of inflamm-aging. Ageing Res Rev 7, 83–105.

Salmon, A.B., Murakami, S., Bartke, A., Kopchick, J., Yasumura, K., and Miller, R.A. (2005). Fibroblast cell lines from young adult mice of long-lived mutant strains are resistant to multiple forms of stress. Am J Physiol Endocrinol Metab 289, E23–29.

Salmon, A.B., Sadighi Akha, A.A., Buffenstein, R., and Miller, R.A. (2008). Fibroblasts from naked mole-rats are resistant to multiple forms of cell injury, but sensitive to peroxide, ultraviolet light, and endoplasmic reticulum stress. J Gerontol A Biol Sci Med Sci 63, 232–241.

Schlachetzki, J.C., Saliba, S.W., and Oliveira, A.C. (2013). Studying neurodegenerative diseases in culture models. Rev Bras Psiquiatr 35 *Suppl 2*, S92–100.

Schuster, E., McElwee, J.J., Tullet, J.M., Doonan, R., Matthijssens, F., Reece-Hoyes, J.S., Hope, I.A., Vanfleteren, J.R., Thornton, J.M., and Gems, D. (2010). DamID in C. elegans reveals longevity-associated targets of DAF-16/FoxO. Mol Syst Biol 6, 399.

Sharma, O.P., and Bhat, T.K. (2009). DPPH antioxidant assay revisited. Food Chem 113, 1202–1205.

Shaw, R.J., Bardeesy, N., Manning, B.D., Lopez, L., Kosmatka, M., DePinho, R.A., and Cantley, L. C., (2004). The LKB1 tumor suppressor negatively regulates mTOR signaling. Cancer Cell 6, 91–99.

Shimokawa, I., Komatsu, T., Hayashi, N., Kim, S.E., Kawata, T., Park, S., Hayashi, H., Yamaza, H., Chiba, T., and Mori, R. (2015). The life-extending effect of dietary restriction requires Foxo3 in mice. Aging Cell 14, 707–709.

Shin, B.Y., Jin, S.H., Cho, I.J., and Ki, S.H. (2012). Nrf2-ARE pathway regulates induction of Sestrin-2 expression. Free Radic Biol Med 53, 834–841.

Singh, A., Boldin-Adamsky, S., Thimmulappa, R.K., Rath, S.K., Ashush, H., Coulter, J., Blackford, A., Goodman, S.N., Bunz, F., Watson, W.H., et al. (2008). RNAi-mediated silencing of nuclear factor erythroid-2-related factor 2 gene expression in non-small cell lung cancer inhibits tumor growth and increases efficacy of chemotherapy. Cancer Res 68, 7975–7984.

Singh, S.P., Niemczyk, M., Saini, D., Sadovov, V., Zimniak, L., and Zimniak, P. (2010). Disruption of the mGsta4 gene increases life span of C57BL mice. J Gerontol A Biol Sci Med Sci 65, 14–23.

Slack, C., Giannakou, M.E., Foley, A., Goss, M., and Partridge, L. (2011). dFOXO-independent effects of reduced insulin-like signaling in Drosophila. Aging Cell 10, 735–748.

Solis, G.M., and Petrascheck, M. (2011). Measuring Caenorhabditis elegans life span in 96 well microtiter plates. J Vis Exp.

Solomon, V.R., and Lee, H. (2012). Anti-breast cancer activity of heteroaryl chalcone derivatives. Biomed Pharmacother 66, 213–220.

Steinbaugh, M.J., Sun, L.Y., Bartke, A., and Miller, R.A. (2012). Activation of genes involved in xenobiotic metabolism is a shared signature of mouse models with extended lifespan. Am J Physiol Endocrinol Metab 303, E488–495.

Suzuki, T., Motohashi, H., and Yamamoto, M. (2013). Toward clinical application of the Keap1-Nrf2 pathway. Trends Pharmacol Sci 34, 340–346.

Sykiotis, G.P., and Bohmann, D. (2008). Keap1/Nrf2 signaling regulates oxidative stress tolerance and lifespan in Drosophila. Dev Cell 14, 76–85.

Tan, N.S., and Wahli, W. (2014). The emerging role of Nrf2 in dermatotoxicology. EMBO Mol Med 6, 431–433.

Thorne, N., Auld, D.S., and Inglese, J. (2010). Apparent activity in high-throughput screening: origins of compound-dependent assay interference. Curr Opin Chem Biol 14, 315–324.

Tullet, J.M., Hertweck, M., An, J.H., Baker, J., Hwang, J.Y., Liu, S., Oliveira, R.P., Baumeister, R., and Blackwell, T.K. (2008). Direct inhibition of the longevity-promoting factor SKN-1 by insulin-like signaling in C. elegans. Cell 132, 1025–1038.

Tusher, V.G., Tibshirani, R., and Chu, G. (2001). Significance analysis of microarrays applied to the ionizing radiation response. Proc Natl Acad Sci U S A 98, 5116–5121.

Walters, H.E., Deneka-Hannemann, S., and Cox, L.S. (2016). Reversal of phenotypes of cellular senescence by pan-mTOR inhibition. Aging (Albany NY) 8, 231–244.

Walters, W.P., and Namchuk, M. (2003). Designing screens: how to make your hits a hit. Nat Rev Drug Discov 2, 259–266.

Wang, T.T., Zeng, G.C., Li, X.C., and Zeng, H.P. (2010). In vitro studies on the antioxidant and protective effect of 2-substituted −8-hydroxyquinoline derivatives against H(2)O(2)-induced oxidative stress in BMSCs. Chem Biol Drug Des 75, 214–222.

Wessells, R.J., Fitzgerald, E., Cypser, J.R., Tatar, M., and Bodmer, R. (2004). Insulin regulation of heart function in aging fruit flies. Nat Genet 36, 1275–1281.

Wilkinson, J.E., Burmeister, L., Brooks, S.V., Chan, C.C., Friedline, S., Harrison, D.E., Hejtmancik, J.F., Nadon, N., Strong, R., Wood, L.K., et al. (2012). Rapamycin slows aging in mice. Aging Cell 11, 675–682.

Wolfson, R.L., Chantranupong, L., Saxton, R.A., Shen, K., Scaria, S.M., Cantor, J.R., and Sabatini, D.M. (2016). Sestrin2 is a leucine sensor for the mTORC1 pathway. Science 351, 43–48.

Yadav, V.R., Prasad, S., Sung, B., and Aggarwal, B.B. (2011). The role of chalcones in suppression of NF-kappaB-mediated inflammation and cancer. Int Immunopharmacol 11, 295–309.

Yang, Y.L., Loh, K.S., Liou, B.Y., Chu, I.H., Kuo, C.J., Chen, H.D., and Chen, C.S. (2013). SESN-1 is a positive regulator of lifespan in Caenorhabditis elegans. Exp Gerontol 48, 371–379.

Ye, X., Linton, J.M., Schork, N.J., Buck, L.B., and Petrascheck, M. (2014). A pharmacological network for lifespan extension in Caenorhabditis elegans. Aging Cell 13, 206–215.

Yeh, E.T. (2009). SUMOylation and De-SUMOylation: wrestling with life’s processes. J Biol Chem 284, 8223–8227.

Zhang, G., Li, J., Purkayastha, S., Tang, Y., Zhang, H., Yin, Y., Li, B., Liu, G., and Cai, D. (2013). Hypothalamic programming of systemic ageing involving IKK-beta, NF-kappaB and GnRH. Nature 497, 211–216.

Zhang, S., Lin, Y., Kim, Y.S., Hande, M.P., Liu, Z.G., and Shen, H.M. (2007). c-Jun N-terminal kinase mediates hydrogen peroxide-induced cell death via sustained poly(ADP-ribose) polymerase-1 activation. Cell Death Differ 14, 1001–1010.

Zhou, W., Ryan, J.J., and Zhou, H. (2004). Global analyses of sumoylated proteins in Saccharomyces cerevisiae. Induction of protein sumoylation by cellular stresses. J Biol Chem 279, 32262–32268.

Zoncu, R., Efeyan, A., and Sabatini, D.M. (2011). mTOR: from growth signal integration to cancer, diabetes and ageing. Nat Rev Mol Cell Biol 12, 21–35.

