## Supplemental Figure S1 to S10 for "Stress Resistance Screen in a Human Primary Cell Line Identifies Small Molecules that Affect Aging Pathways and Extend *C. elegans’* Lifespan"

**Figure S1. Increased oxidative-stress resistance upon *AKT1* or *KEAP1* knockdown (related to Figure 1)**

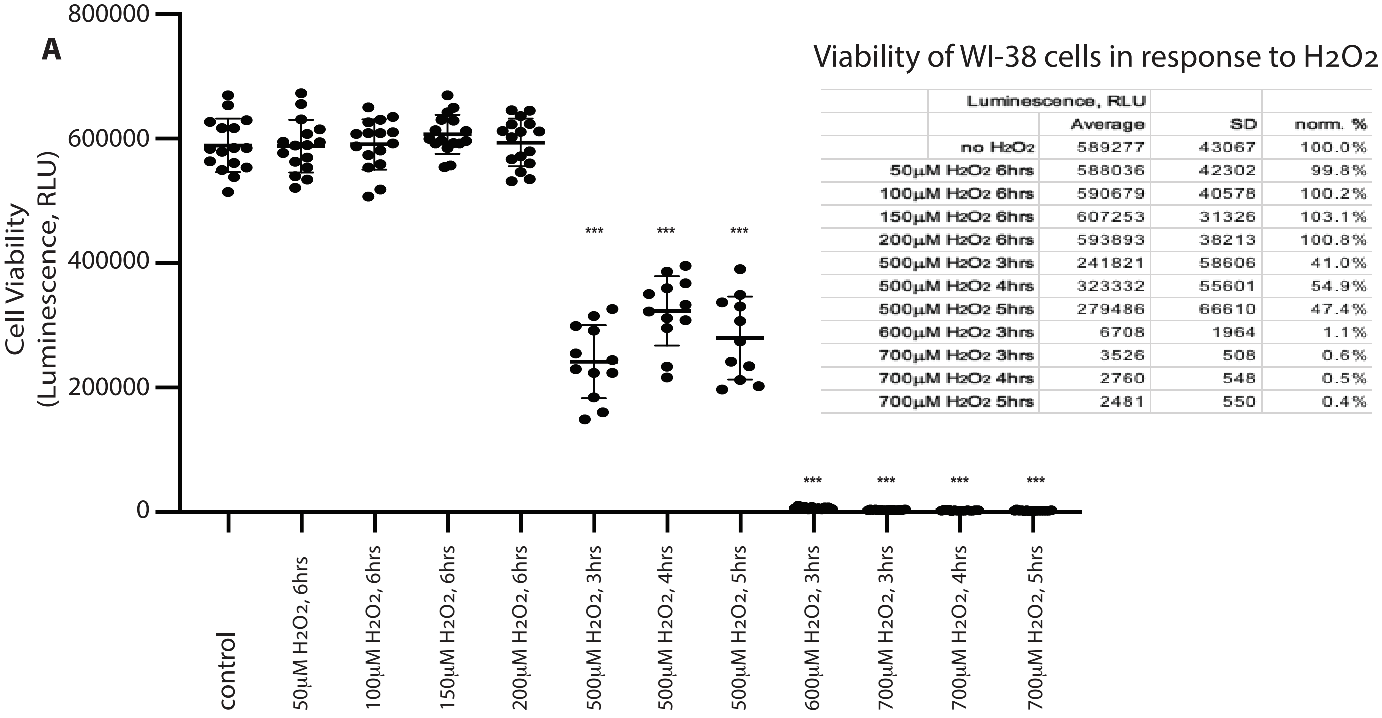


Viability of WI-38 cells in response to $H_2O_2$

| | Luminescence, RLU | | |
|---|---|---|---|
| | Average | SD | norm. % |
| no $H_2O_2$ | 589277 | 43067 | 100.0% |
| 50μM $H_2O_2$ 6hrs | 588036 | 42302 | 99.8% |
| 100μM $H_2O_2$ 6hrs | 590679 | 40578 | 100.2% |
| 150μM $H_2O_2$ 6hrs | 607253 | 31326 | 103.1% |
| 200μM $H_2O_2$ 6hrs | 593893 | 38213 | 100.8% |
| 500μM $H_2O_2$ 3hrs | 241821 | 58606 | 41.0% |
| 500μM $H_2O_2$ 4hrs | 323332 | 55601 | 54.9% |
| 500μM $H_2O_2$ 5hrs | 279486 | 66610 | 47.4% |
| 600μM $H_2O_2$ 3hrs | 6708 | 1964 | 1.1% |
| 700μM $H_2O_2$ 3hrs | 3526 | 508 | 0.6% |
| 700μM $H_2O_2$ 4hrs | 2760 | 548 | 0.5% |
| 700μM $H_2O_2$ 5hrs | 2481 | 550 | 0.4% |

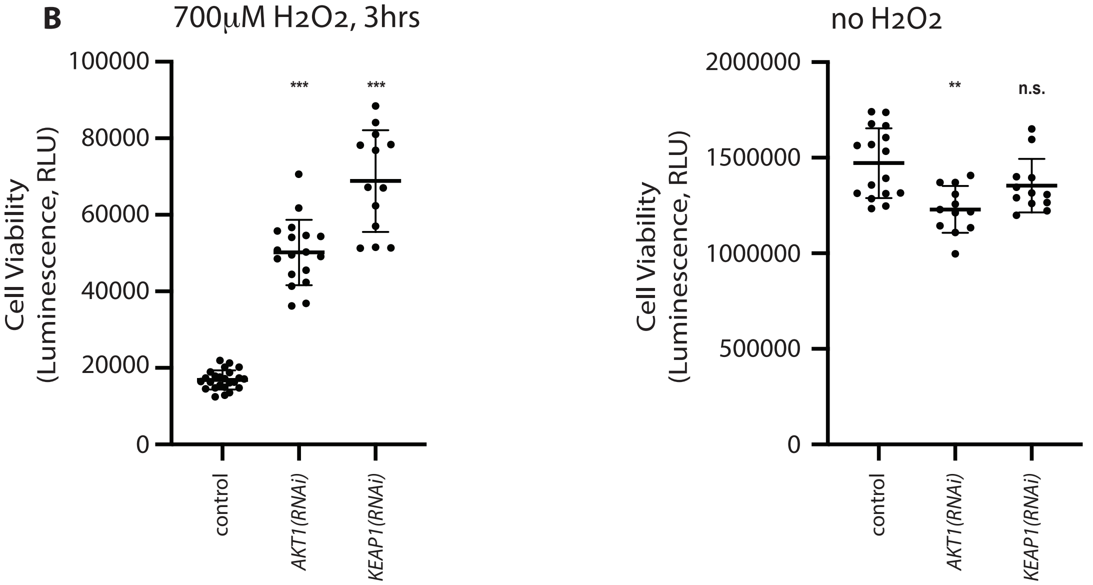


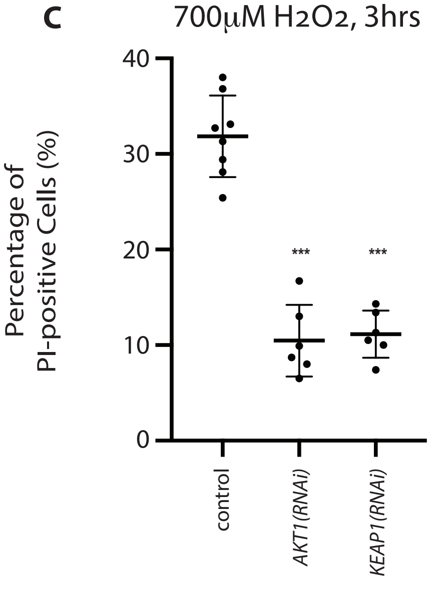

**Figure S2. Z-prime scores across the screen (related to Figure 1)**

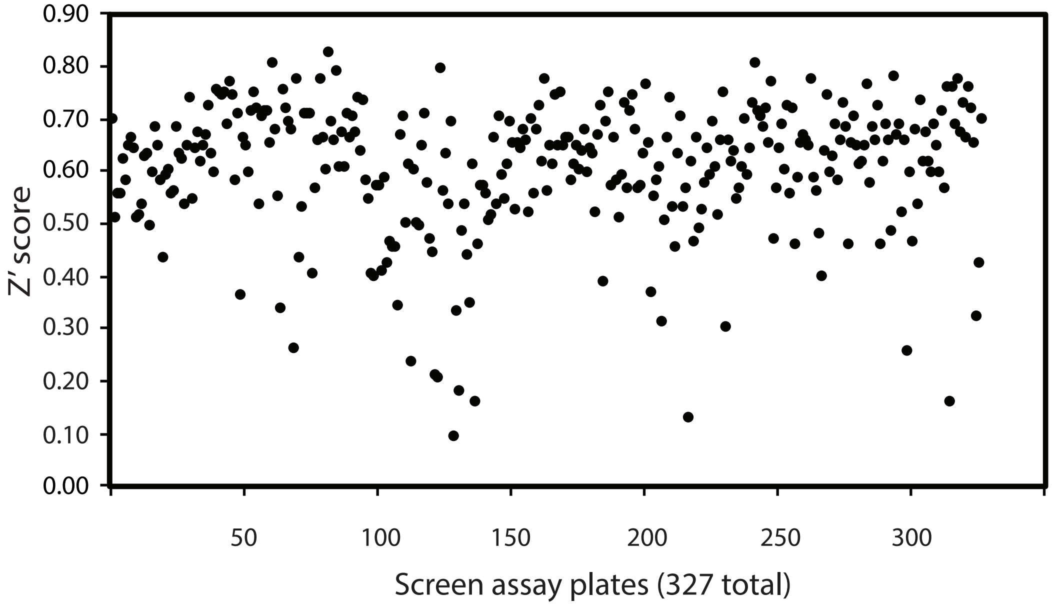


**Figure S3. ROS-scavenging capacity of certain small molecules in the absence of cells (related to Figure 1)**

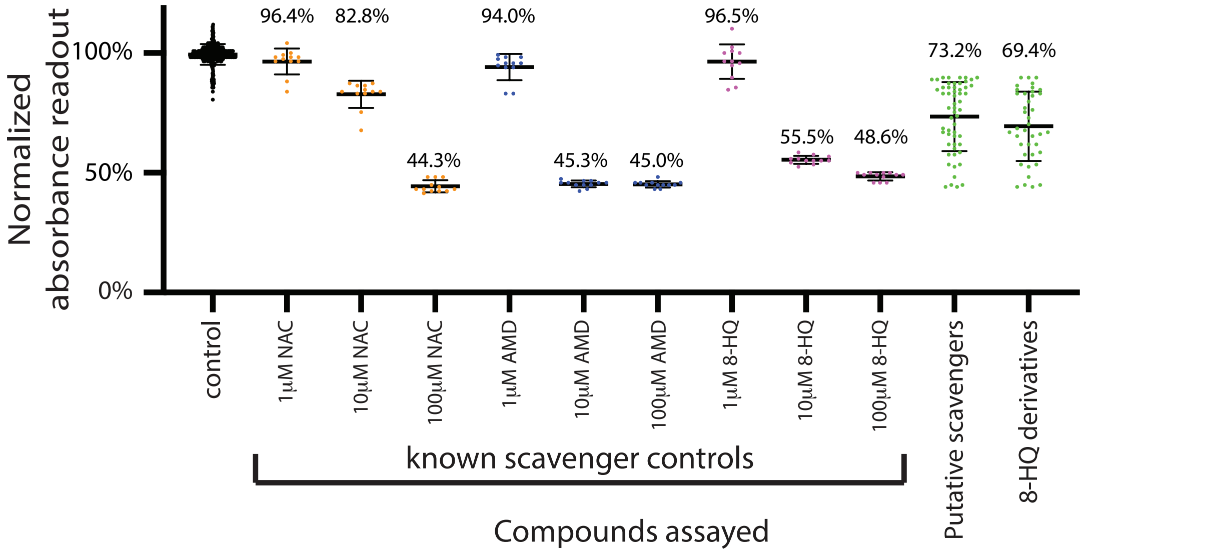


**Figure S4. No H2O2-quenching effects by small molecules in the absence of cells (related to Figure 2)**

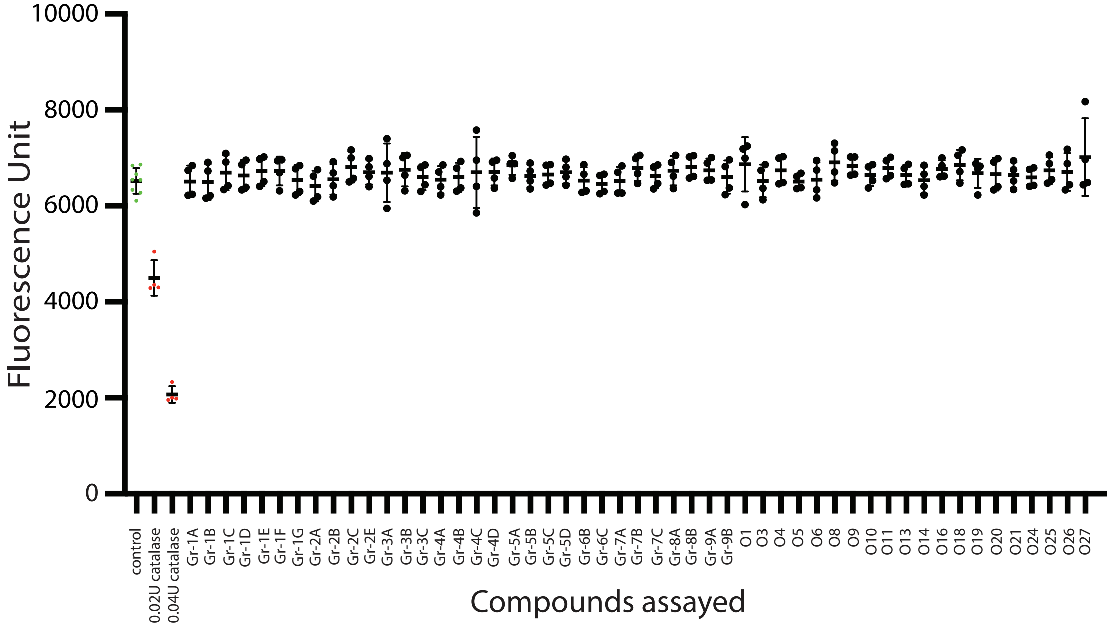

**Figure S5-1. Long-term effects of small molecules on ATP levels (related to Figure 3)**

**A**

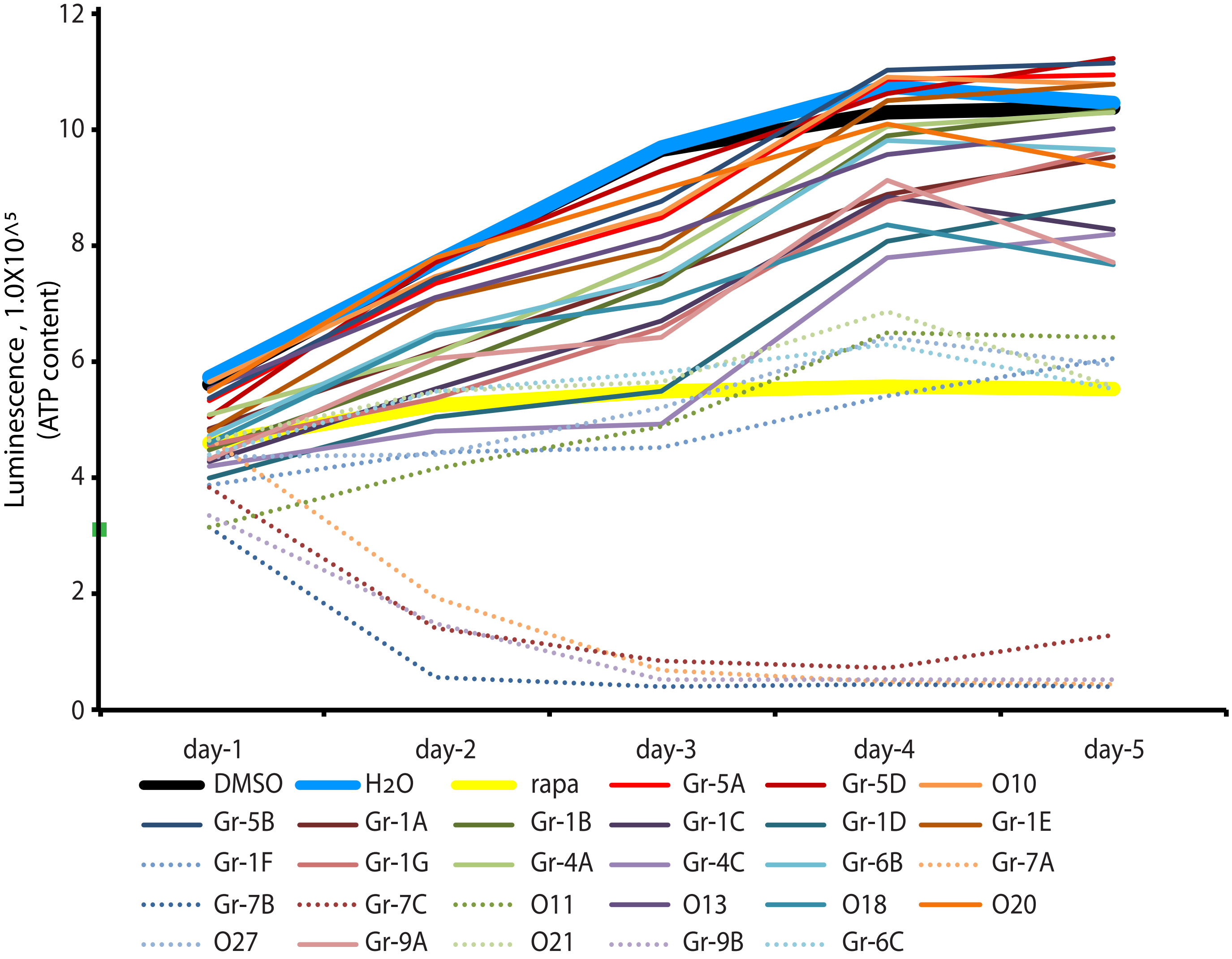


**B**

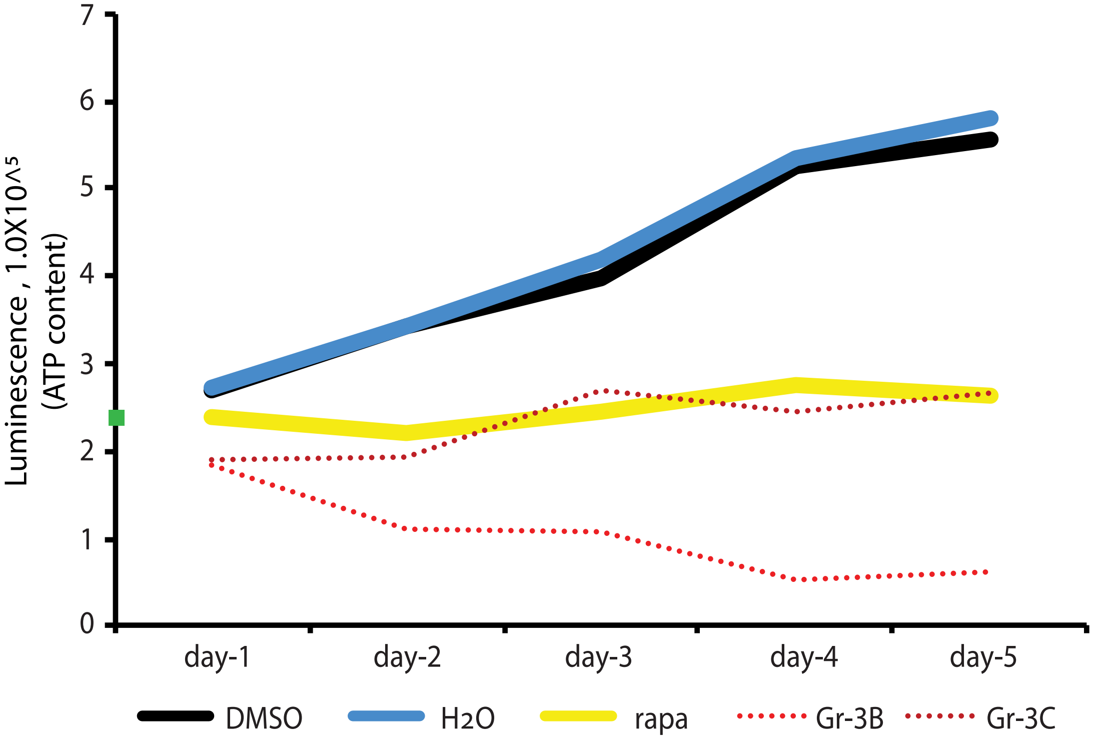

**Figure S5-2. Effects of prolonged small-molecule incubation on WI-38 cell confluency (related to Figure 3)**

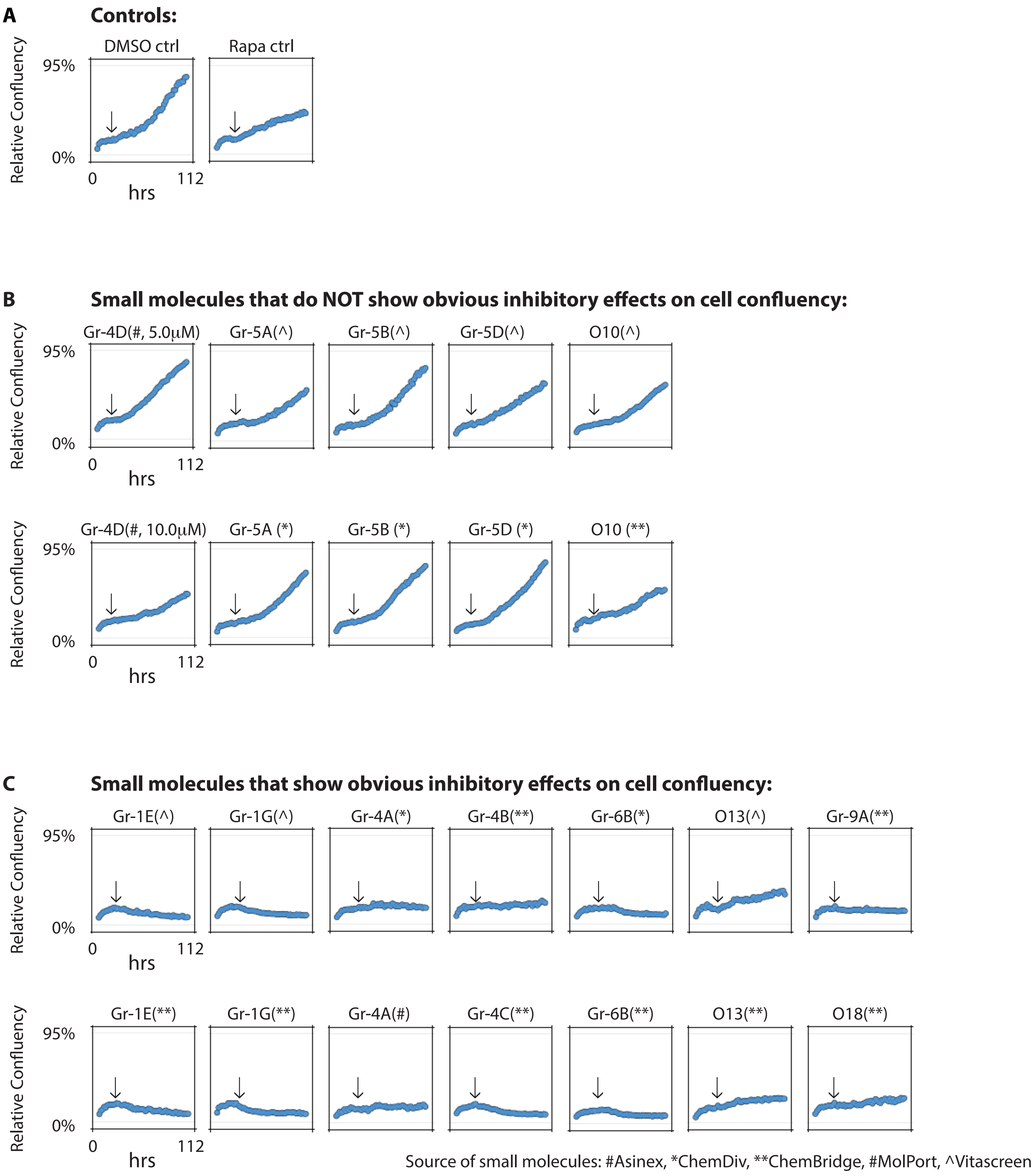

### Figure S6. Increased cells viability upon $H_2O_2$ by small molecules (related to Figure 3)

A

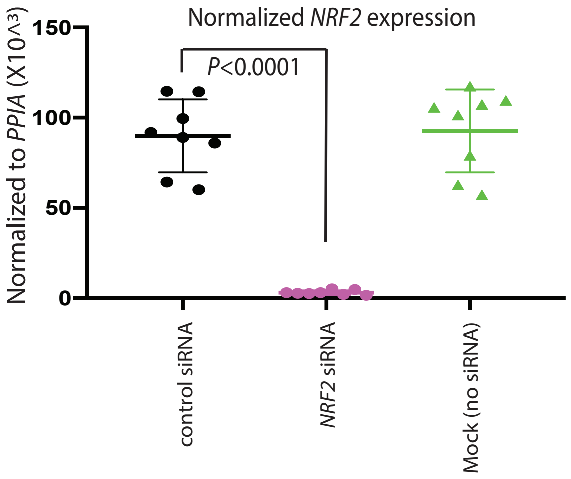


B

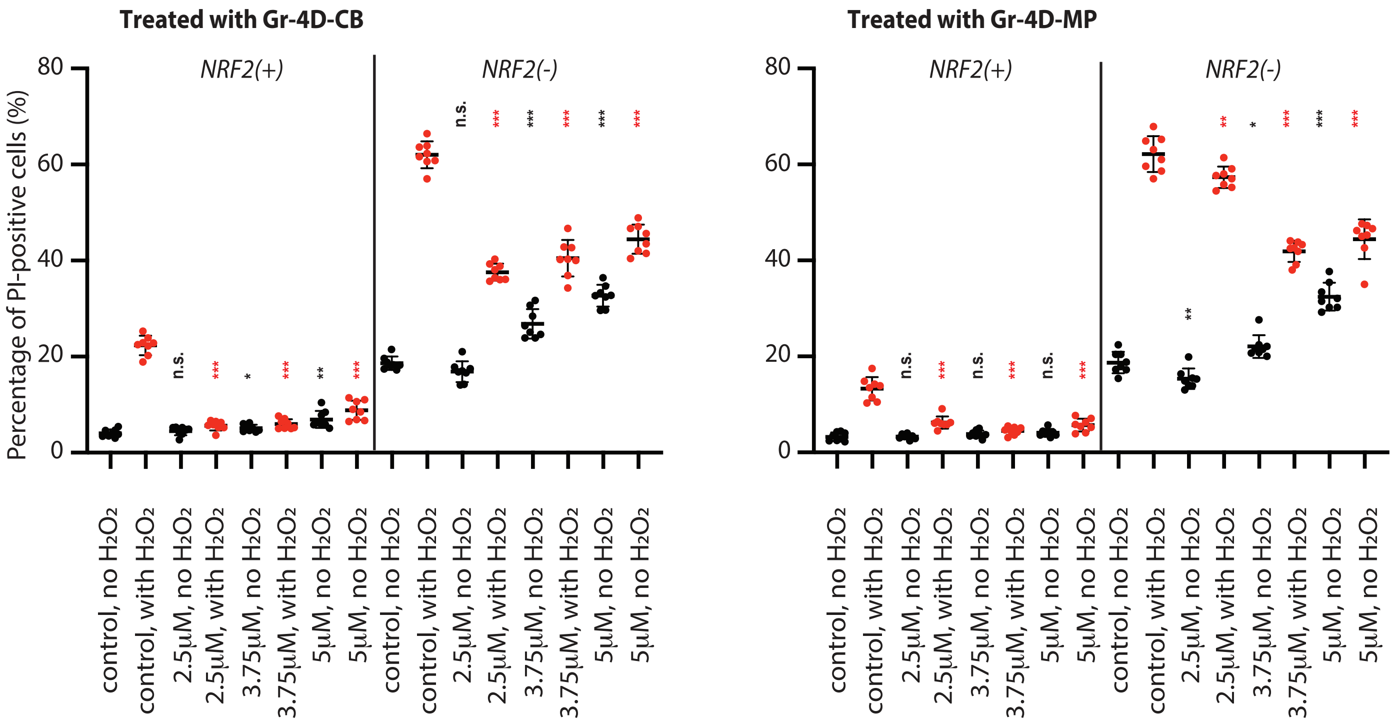


C

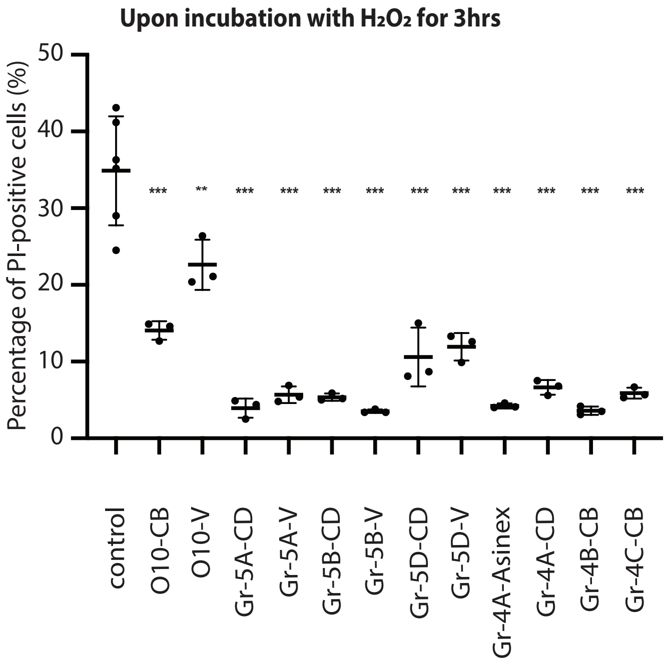

**Figure S7. Effects of small molecules on gene expression of WI-38 cells (related to Figure 3)**

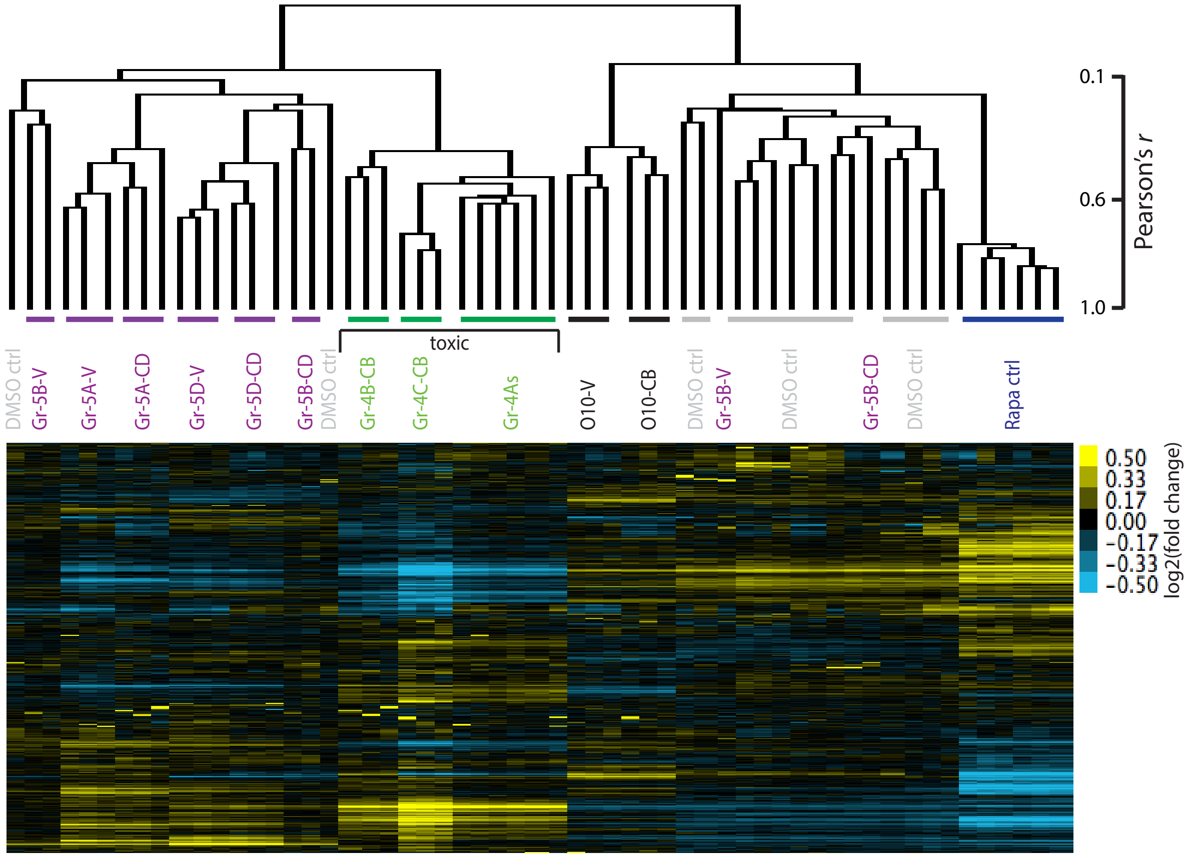

**Figure S8. Inhibitory effects of certain small molecules on PARP (related to Figure 2)**

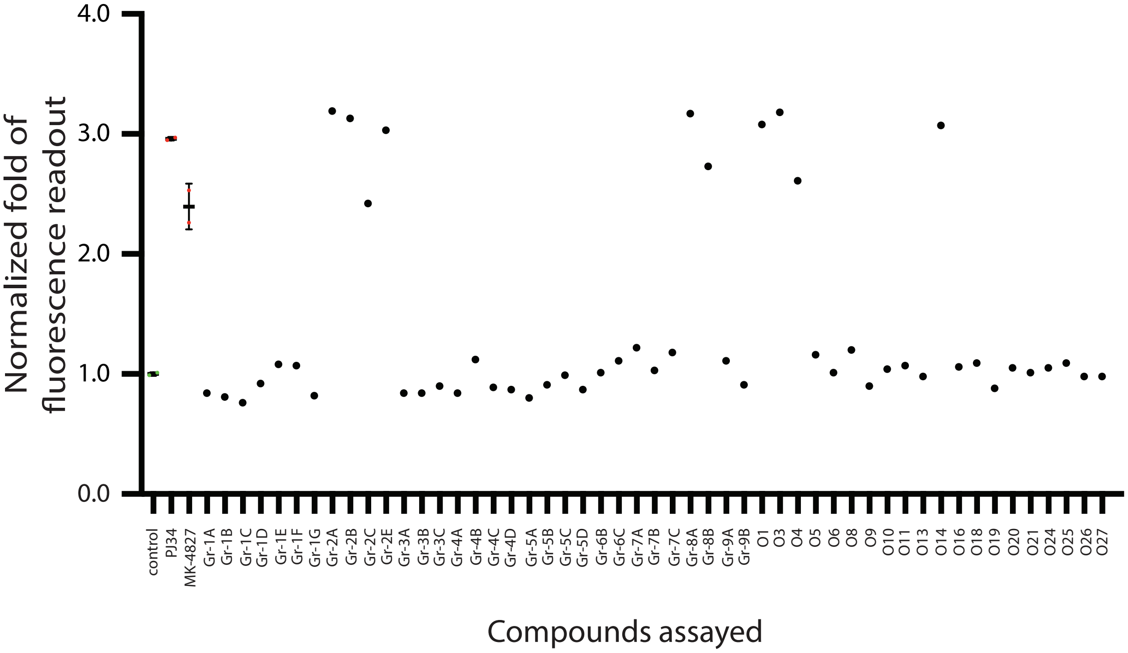

**Figure S9. Protective effects of certain small molecules against poly-Q toxicity (related to Table 1)**

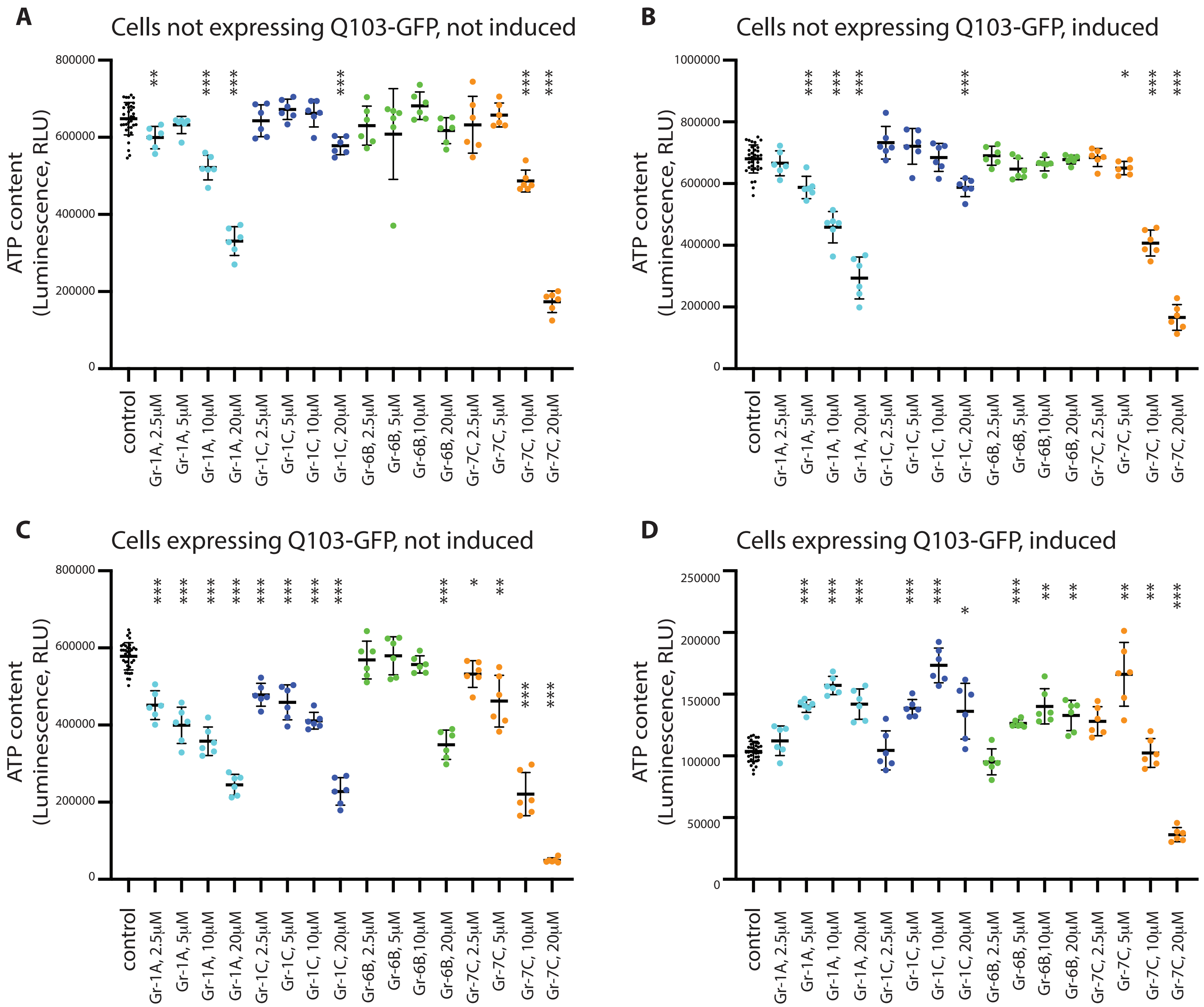

**Figure S10. Effects of Gr-4D on *C. elegans* expressing several pathway reporters (related to Figure 4).**

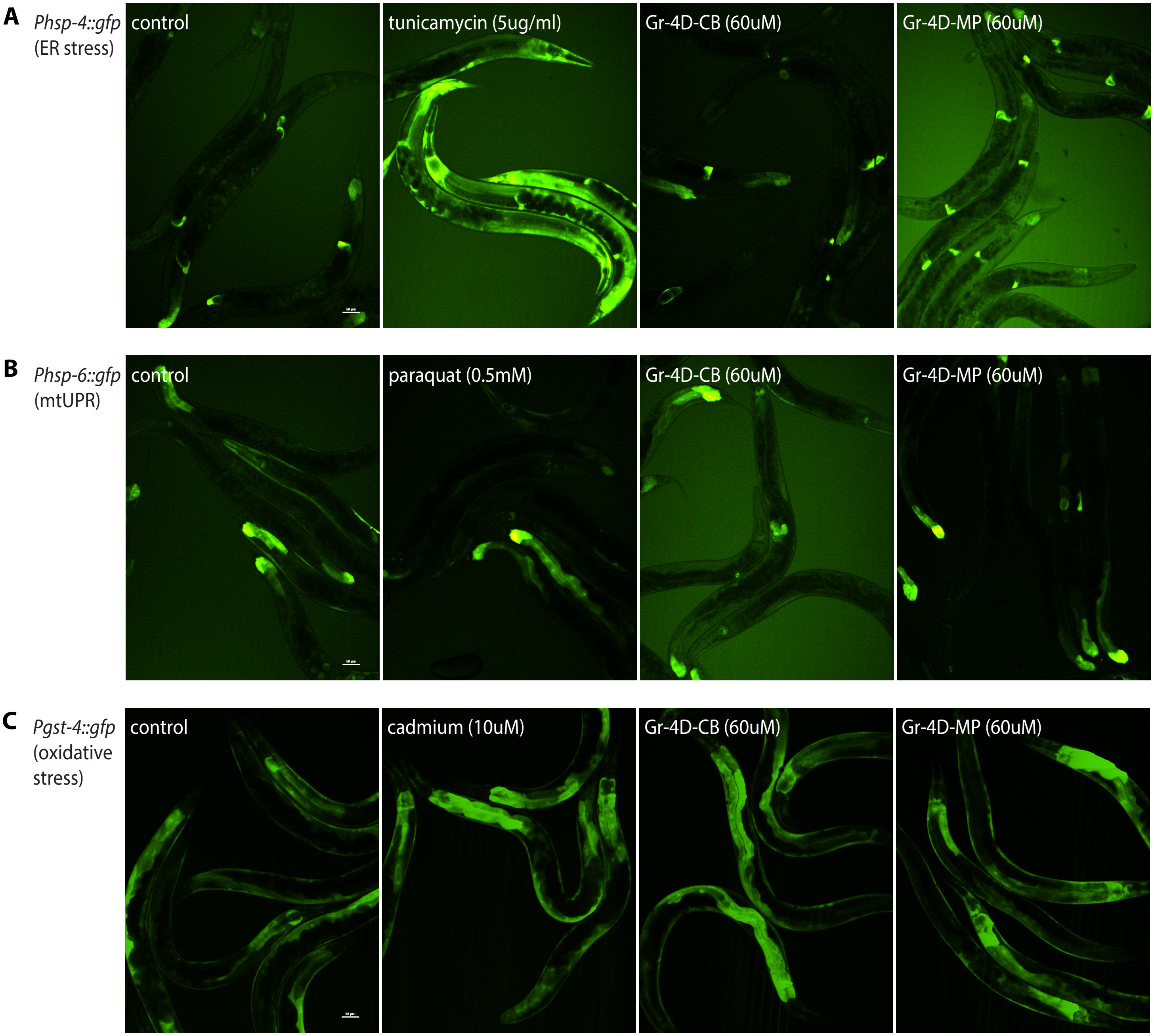

**Supplemental Figure Legends**

Supplemental Figure 1 (related to Figure 1). Increased oxidative-stress resistance upon *AKT1* or *KEAP1* knockdown.

A) To define the optimal assay conditions for our screen, WI-38 cells were incubated with different doses of $H_2O_2$ for different periods of time in 384-well plates and then assayed for viability by measuring the ATP content. Shown here are representative data, indicating that high doses (above 600 $\mu$M) of $H_2O_2$ substantially depleted the ATP content after 3 hours of incubation (luminescence values are shown by the side). Control, n = 16; for all other $H_2O_2$ test samples, n = 11-16 (***, $P$ <0.0001 for 500 $\mu$M $H_2O_2$ and above, Student's t-test). Shown in all the figures, unless otherwise specified, were mean ± SD.

B) WI-38 cells were transfected with 20 nM AllStars negative control or *AKT1* or *KEAP1* siRNA oligos (Qiagen) and then assayed for viability 72 hours post-transfection upon $H_2O_2$. Representative data indicated that *AKT1* or *KEAP1* knockdown significantly increased cell viability, as measured by ATP content following 3 hours incubation in 700 $\mu$M $H_2O_2$. One-way ANOVA was performed for statistics ($p<0.001$), plus post-comparison with control (***, $p<0.001$). Control, n = 24; *AKT1* knockdown, n = 18; *KEAP1* knockdown, n = 13. Viability was also measured in the same experiment for cells not subjected to $H_2O_2$. One-way ANOVA ($p<0.001$), plus post-comparison with control (**, $p<0.01$; n.s., not significant). Control, n = 16; *AKT1* knockdown, n = 12; *KEAP1* knockdown, n = 12.

C) Propidium iodide staining was performed in parallel to analyze the fraction of dead/dying cells upon $H_2O_2$. Consistent with the ATP assay results, *AKT1* or *KEAP1* knockdown significantly reduced the fraction of PI-positive cells. One-way ANOVA ($p<0.001$), plus post-comparison with control (***, $p<0.001$). Control, n = 8; *AKT1* knockdown, n = 6; *KEAP1* knockdown, n = 6.
